# Rewiring *Escherichia coli* to Transform Formate into Methyl Groups

**DOI:** 10.1101/2024.11.27.625645

**Authors:** Michael K. F. Mohr, Ari Satanowski, Steffen N. Lindner, Tobias J. Erb, Jennifer N. Andexer

## Abstract

**Background:** *Biotechnological applications are steadily growing and have become an important tool to reinvent the synthesis of chemicals and pharmaceuticals for lower dependence on fossil resources. In order to sustain this progression, new feedstocks for biotechnological hosts have to be explored. One-carbon (C_1_-)compounds, including formate, derived from CO_2_ or organic waste are accessible in large quantities with renewable energy, making them promising candidates. Previous studies showed that introduction of the formate assimilation machinery from* Methylorubrum extorquens *into* Escherichia coli *allows assimilation of formate into the established biotechnological host. Applying this established route for formate assimilation, we here investigated utilisation of formate for the production of value-added building blocks in* E. coli using S*-adenosylmethionine (SAM)-dependent methyltransferases*.

**Results:** *We first analysed methylation activity in* E. coli *BL21* with a two-vector system to produce three different methyltransferases together with the formate assimilation machinery. *Feeding isotopically labelled formate, products with 51 – 81% ^13^C-labelling could be obtained by maintaining* in vivo *methylation activity. Focussing on improvement of* in vivo *methylation, we analysed two further* E. coli *strains with an engineered C_1_-metabolism and, following condition optimisation, achieved a doubled methylation activity with a share of more than 70% formate-derived methyl groups*.

**Conclusions:** *This study demonstrates the efficient transformation of formate into methyl groups in* E. coli*. Our findings support that feeding formate can improve the availability of usable C_1_-compounds and, as a result, increase* in vivo *methylation activity in engineered* E. coli.

## Background

The majority of industrial processes depends on fossil resources for the synthesis of materials such as plastics, chemicals, or pharmaceuticals.^[1,2]^ With depleting fossil resources and the emerging effects of climate change, defossilisation is an important strategy to create a greener and more sustainable chemical industry.^[1]^ Development of biochemical applications utilising alternative and renewable feedstocks to support or replace existing processes is key for this transformation.^[3,4]^ A promising approach is the application of CO_2_-derived feedstocks in chemobiohybrid processes, in which the greenhouse gas is first (electro)chemically converted into reduced one-carbon (C_1_-)compounds, which are further valorised through biotechnological efforts.^[5–7]^ Formate is such a compound with the benefit of stability, biodegradability, ease of handling and low toxicity.^[7,8]^ For this reason, the development of biotechnological strategies based on the C_1_-compound formate is an emerging field.^[7–9]^

In nature, one important strategy to transfer C_1_-compounds onto various molecules is *S*-adenosylmethionine (SAM)-dependent methylation catalysed by methyltransferases (MTs, Figure 1 **a**).^[10–12]^ Many natural products with biological activity are methylated and methylation of pharmaceuticals at specific positions has been shown to increase their potency by up to three orders of magnitude, called the “magic methyl effect”.^[13]^ This makes selective methylation an important step in lead structure optimisation during development of new pharmaceuticals (Figure 1 **b**). Chemical introduction of methyl groups is in many cases challenging to direct and commonly mediated by methylation agents such as methyl iodide or diazomethane which are mutagenic and toxic to the environment.^[14]^ In comparison to chemical methylation, MTs introduce the methyl group stereo-, regio- and chemoselectively, only synthesising one specific product, and by investigating radical SAM MTs, chemically inaccessible sp^3^-hybridised carbons could be targeted as well.^[10,15]^ SAM-dependent methylation using biotechnological organisms such as *E. coli* could allow replacement of methylation agents in combination with fewer byproduct formation for many applications.^[11,16,17]^ *In vivo* methylation activity is often a limiting factor when natural compounds are produced in biotechnological hosts, emphasising the need to develop strategies to improve *in vivo* methylation activity.^[16–19]^

**Figure 1.**
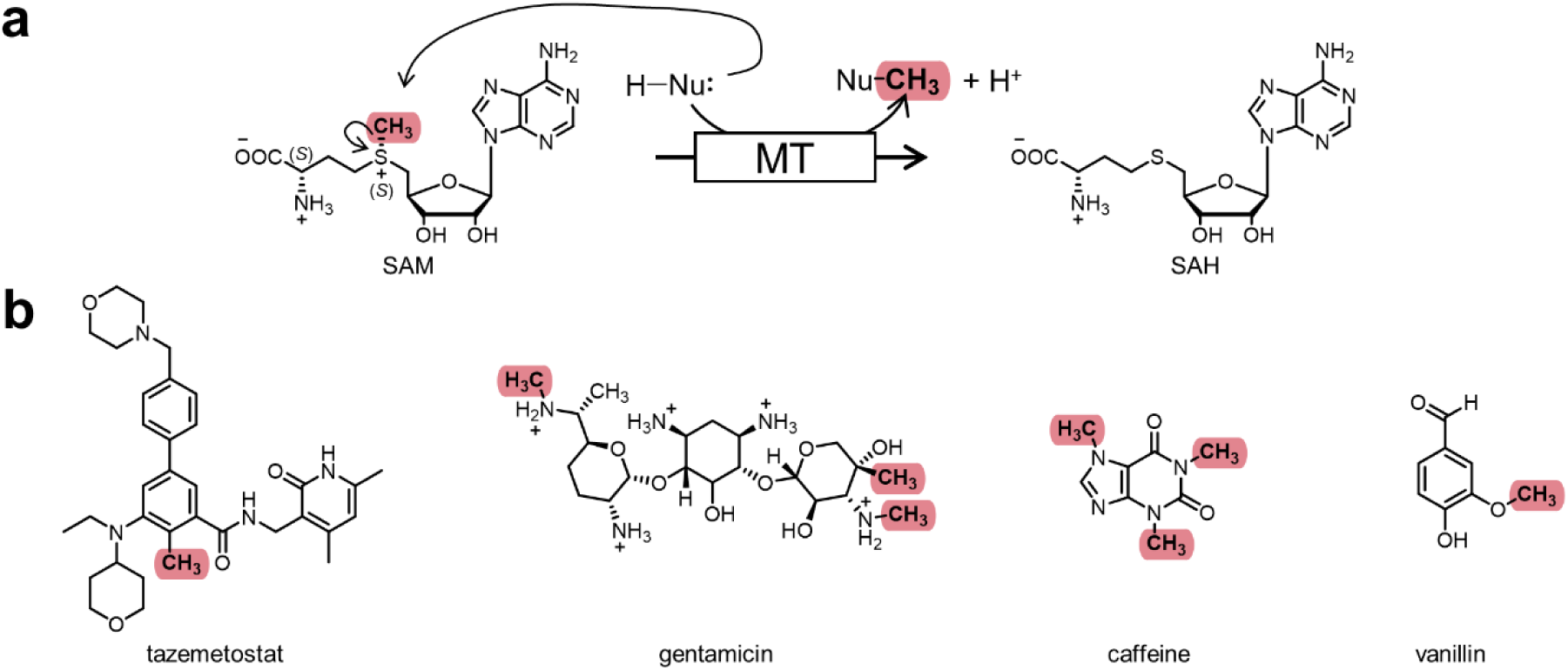
SAM-dependent methylation. **a** Reaction mechanism of SAM-dependent methylation by MTs. **b** Pharmaceuticals and natural compounds carrying magic methyl groups, significantly influencing potency. *C-*methylation of gentamicin is mediated by a radical SAM MT, other methyl groups are introduced by conventional MTs. SAH = *S*-adenosylhomocysteine. *E. coli*.^[9,28,32]^ Reduction of methylene-H_4_F by the NADH-dependent methylene-H_4_F reductase (*Ec*MetF, Figure 2) yields methyl-H_4_F.^[33]^ *Ec*MetE or *Ec*MetH transfer the methyl group onto homocysteine for production of the proteinogenic amino acid methionine, the precursor of SAM.^[23]^ Introduction of FCM in engineered *E. coli* strains allowed all methyl groups of methionine to derive from formate when ^13^C-labelled formate was fed, demonstrating the potential to use formate for the synthesis of methyl groups.^[34]^ Growth of the synthetic formatotrophs solely on formate is approximately 10-times slower than on established feedstocks,^[9,32]^ making a strategy with growth from established feedstocks and supply of formate to push methylation of selected building blocks a great option.

Optimisation of intracellular SAM availability and acceleration of the cofactor’s regeneration can improve *in vivo* methylation activities and, therefore, increase the yield of desired methylated products.^[16–20]^ One strategy which proved successful is increasing concentrations of the SAM precursor methionine and its regeneration from homocysteine.^[17,20–22]^ In *E. coli*, homocysteine is methylated to methionine through transfer of a methyl group from methyl-tetrahydrofolate (methyl-H_4_F) catalysed by the methionine synthases *Ec*MetE and *Ec*MetH (Figure 2).^[23]^ In *E. coli’*s native metabolism, the C_1_-H_4_F pool is fuelled from serine and glycine, catalysed by serine hydroxymethyltransferase (*Ec*GlyA) and the glycine cleavage system (*Ec*GCV), respectively.^[24,25]^ The C_1_-H_4_F pool supplies C_1_-carbon units for the synthesis of nucleobases and amino acids such as methionine or histidine as well as for formylation of methionine, making it essential for protein production and cell growth.^[6,26,27]^ In a study by *Okano et al.,* it was shown that increasing the availability of C_1_-compounds to improve methyl-H_4_F production benefits *in vivo* methylation activity in *E. coli*.^[22]^ The C_1_-compound formate could be an ideal supplier of C_1_-building blocks to fuel the C_1_-H_4_F metabolism of biotechnological hosts and drive *in vivo* methylation forward.

**Figure 2.**
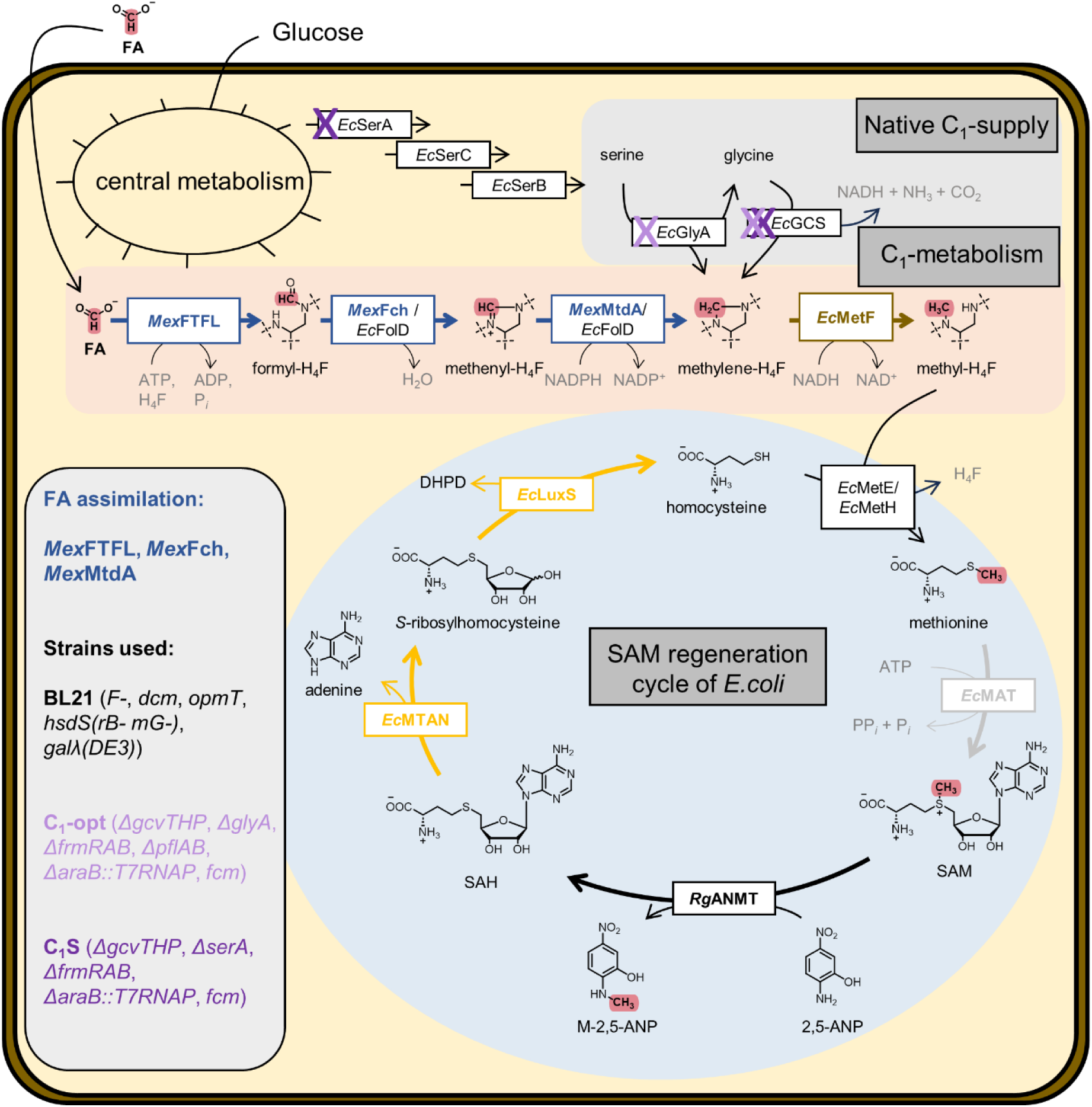
Metabolic network for *in vivo* methylation from formate. Formate assimilation with *Mex*FTFL, *Mex*Fch and *Mex*Mtda (together abbreviated as “FCM”) is shown embedded in the native metabolism of *E. coli* in combination with the given C_1_-building block supply pathways, the C_1_-H_4_F metabolism and the SAM regeneration cycle. Strains investigated in this study are listed and genes deleted are labelled in the respective colour.

Within previous publications, strategies to assimilate formate into the metabolism of *E. coli* have been established.^[9,28]^ Introduction of the formate-H_4_F ligase from *Methylorubrum extorquens* (*Mex*FTFL) into *E. coli* allows synthesis of formyl-H_4_F from formate and H_4_F in an adenosine-5’-triphosphate (ATP)-dependent reaction (Figure 2).^[9,29,30]^ Further, introduction of formyl-H_4_F cyclohydrolase (*Mex*Fch) and methylene-H_4_F dehydrogenase (*Mex*MtdA), both from *M. extorquens*, cyclise and reduce formyl-H_4_F to methylene-H_4_F, the central metabolite of the C_1_-H_4_F metabolism.^[29–31]^ This formate assimilation strategy composed of *Mex*FTFL, *Mex*Fch and *Mex*MtdA (together referred to as FCM) proved powerful and was used for full assembly of all carbon scaffolds from formate and CO_2_ in synthetic formatotrophic

In order to activate the methyl group, native methionine adenosyltransferases (*Ec*MAT, Figure 2) convert ATP and methionine to SAM as a first step of the SAM regeneration cycle.^[35]^ This cofactor is subsequently utilised by MTs for methylation of their substrate, thereby introducing the C_1_-compound into the product. Byproduct of the methylation reaction is *S*-adenosylhomocysteine (SAH), a known inhibitor of MTs.^[10]^ In native metabolism, SAH is degraded in an irreversible reaction to adenine and *S*-ribosylhomocysteine by methyl thioadenosine/SAH nucleosidase (*Ec*MTAN).^[36]^ *S*-ribosylhomocysteine is cleaved by *S*-ribosylhomocysteine lyase (*Ec*LuxS) to homocysteine and the quorum sensing precursor 4,5-dihydroxypentane-2,3-dione (DHPD).^[37]^ Another option to degrade SAH are SAH hydrolases (SAHH) that cleave SAH to adenosine and homocysteine.^[38]^ Transfer of a methyl group from methyl-H_4_F regenerates methionine from homocysteine allowing repetition of methylation with another C_1_-building block.

With this work, we set out to investigate formate as a supplementary source of cellular C_1_-units to ramp up *in vivo* methylation activity in *E. coli*. We first analysed application of formate assimilation through FCM to incorporate the renewable C_1_-compound into methylated products and continued building on these results to improve *in vivo* methylation activity by increasing the availability of C_1_-compounds to fuel the C_1_-H_4_F metabolism.

## Results

As starting point to analyse the influence of formate supplementation on *in vivo* methylation, we used our previously reported model reaction: Anthranilate *N*-MT from *Ruta graveolens* (*Rg*ANMT) transfers a methyl group in a SAM-dependent reaction onto the amino group of the unnatural substrate 2,5-amino nitrophenol (2,5-ANP) producing *N*-methyl-2,5-ANP (M-2,5-ANP, Figure 2).^[20,39]^ The unnatural substrate and product undergo minimal degradation in *E. coli* and can freely permeate in and out of the bacterial cell, making this system a well-suited model for investigations on *in vivo* methylation activity. *In vivo* methylation activity of this model MT-substrate system was shown to be strongly dependent on the availability of methionine, resulting in a great potential to use it to explore the influence of formate on *in vivo* methylation activity.^[20]^

We started by transforming *E. coli* BL21 (Gold)DE3 cells (abbreviated as BL21 in the following) with a plasmid carrying *fcm* under the control of a constitutive promoter (pFCM) and a pET28a vector carrying *rganmt*, resulting in BL21*-fcm-rganmt* (Table SI 1 & Table SI 6). To test *in vivo* methylation activity, a biotransformation approach was chosen, where the MT is first overproduced together with FCM in rich medium (lysogeny broth, LB) during the overexpression phase, followed by harvest, and resuspension in glucose containing minimal medium (M9) to normalise the cell densities (final OD_600_ = 3.0) and allow energy regeneration. Subsequently, the biotransformation phase is initiated by adding the methyl acceptor substrate (2,5-ANP) for the *in vivo* methylation reaction together with formate.^[40,41]^

Comparison of BL21*-fcm-rganmt* to a strain without formate assimilation machinery (BL21-*rganmt*) showed that introducing pFCM already improves total methylation activity without addition of formate (Figure 3). A similar behaviour was observed when an empty spectinomycin resistance carrying pCDF vector was co-overexpressed with *mt* genes on a pET28a vector in BL21 in a previous study.^[20]^ This increase was suggested to be caused through introduction of the spectinomycin resistance encoded on pFCM, an adenylyltransferase.^[20]^ Hereby enhanced ATP regeneration increases ATP availability for SAM synthesis and, hence, improves *in vivo* methylation activity.^[20]^ Supplementation with 5 mM ^13^C-formate did not improve *in vivo* methylation activity further, in either of the strains .Analysis with LC-MS/MS, however, revealed that 67% of the product M-2,5-ANP were ^13^C-labelled (Figure 3, Figure SI 1). This incorporation of formate-derived methyl groups did not take place in BL21-*rganmt* highlighting the necessity of pFCM. These first results demonstrated easy conversion of formate into transferable methyl groups by *in vivo* methylation in *E. coli*.

**Figure 3.**
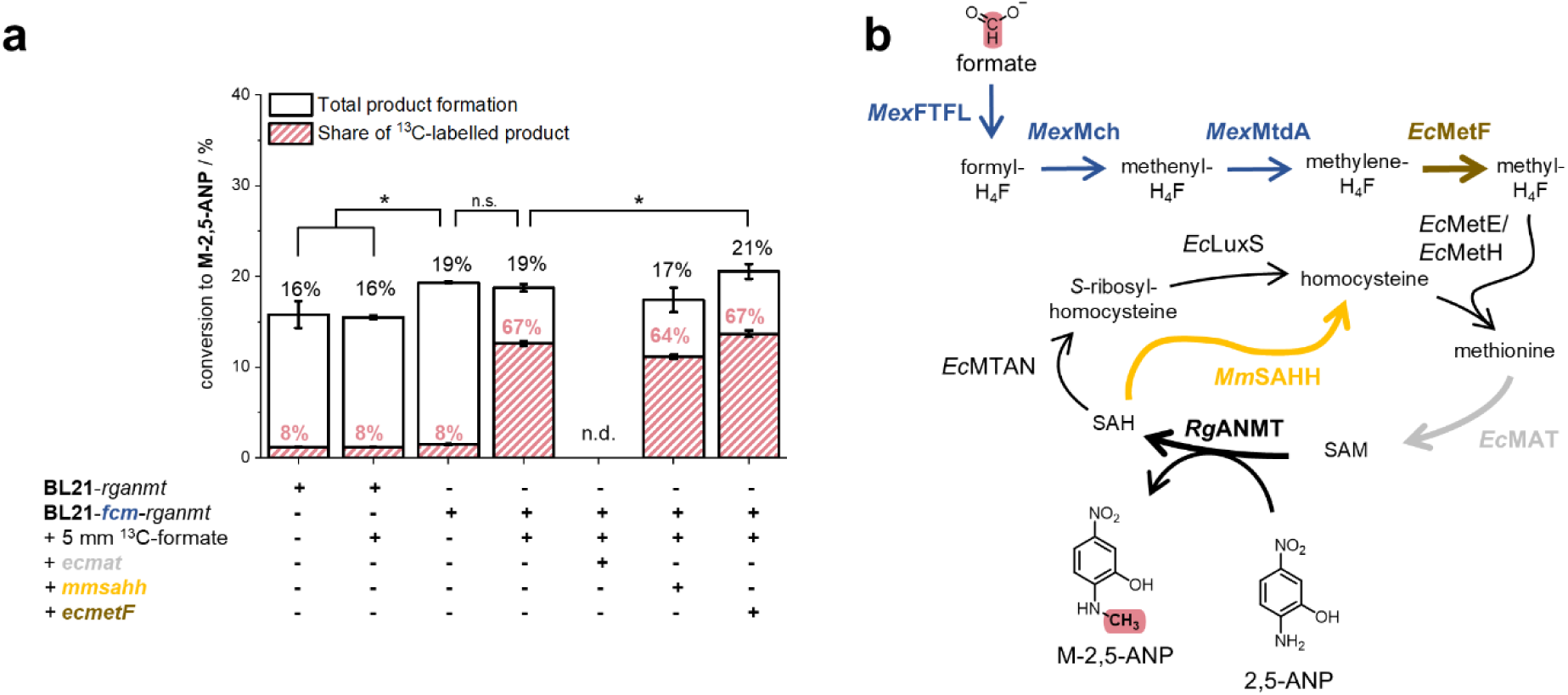
*In vivo* methylation with ^13^C-formate in BL21. **a** Given is the conversion of 2,5-ANP to M-2,5-ANP (black) and the share of ^13^C-labelled product ± SD (red) under different conditions and co-overexpression of genes involved in formate to methyl group transformation or SAM regeneration. **b** Scheme of the metabolic pathway from formate to the methylated product. Experiments were performed in biological triplicates. * = p < 0.05, n.d. = not detected. Experiments were conducted in M9-medium (22 m_M_ glucose) with a final OD_600_ of 3.0, 0.75 m_M_ 2,5-ANP and 0 - 5 m_M_ ^13^C-formate at 37 °C, 170 rpm for 24 h. NOTE: About 8% of M-2,5-ANP is naturally ^13^C-labelled, therefore respective shares are indicators not absolutes for formate-derived methyl groups.

Despite an increased availability of C_1_-building blocks in BL21*-fcm-rganmt* fed with formate, *in vivo* methylation activity did not improve as it was reported in literature.^[22]^ Due to a strong increase in methylation activity when methionine was supplied, neither substrate permeation into the bacterial cell, nor activity of *Rg*ANMT seemed to be the major limiting factors. We rather suspected a kinetic bottleneck either in the SAM regeneration cycle or in the reduction of methylene-H_4_F to methyl-H_4_F through *Ec*MetF. We co-overexpressed *ecmat* and an SAH hydrolase from *Mus musculus* (*mmsahh*), which should improve SAH degradation through cleavage to adenosine and homocysteine; as well as *ecmetf* together with *rganmt* on a pET28a vector. Curiously, BL21-*fcm*-*rganmt*-*ecmat* did not show any methylation activity. An increased metabolic burden could be responsible for the loss of methylation activity of this strain. Co-overexpression of *mmsahh* in BL21-*fcm*-*rganmt*-*mmsahh* lowered methylation activity slightly in combination with a decreased share of labelled methylated product. A 10% increase in methylation activity compared to BL21-*fcm*-*rganmt* supplemented with 5 mM ^13^C-formate was observed with BL21-*fcm*-*rganmt-ecmetf* (Figure 3). Reduction of methylene-H_4_F to methyl-H_4_F by *Ec*MetF, hence, seems to be a minor bottleneck for *in vivo* methylation from formate, even though the increase in methylation activity was rather small, which is why we omitted co-overexpression in the following experiments.

After successful implementation of methylation from formate-derived SAM using *Rg*ANMT in BL21*-fcm-rganmt*, we checked the transferability of this approach to other MTs. The caffeic acid *O*-MT from *Prunus persica* (*Pp*CaOMT) also accepts 2,5-ANP, however catalyses an *O*-methylation to form 2,4-methoxy nitroaniline (2,4-MNA, Figure SI 2). As a third model reaction, the *C*-MT from *Streptomyces rishiriensis* (CouO) was chosen for the transfer of a methyl group onto the unnatural substrate 2,7-dihydroxynaphtalene (DHN), producing 1-methyl-DHN (M-DHN, Figure SI 2). After *in vivo* methylation with supply of 5 mM ^13^C-formate, shares of ^13^C-labelled 2,4-MNA and M-DHN were 51% and 81%, respectively without a substantial change in conversion rates (Figure SI 2). In a previous study, co-overproduction of *Ec*MAT and CouO was found to strongly increase *in vivo* methylation activity; this was also the case here: Co-overproduction of *Ec*MAT together with FCM and CouO increased methylation activity about 3.0-fold with and without the addition of formate. When 5 mM ^13^C-formate were supplied, a share of 68% of M-DHN was ^13^C-labelled (Figure SI 2).

These results demonstrate straight-forward transformation of formate into methyl groups, using BL21-*fcm* in combination with MTs. However, methylation activity did not improve by addition of formate. Initially, we suspected regulation through the transcription repressor *Ec*MetJ to hamper reduction of formate to methyl groups by decreasing the expression of *ecmetF*, *ecmetE*, *ecmetH*, *ecmat* and genes involved in the synthesis of homocysteine.^[42,43]^ Experiments with BL21*ΔmetJ*-*fcm* in combination with the MTs did not result in an increase of methylation activity when supplemented with formate (Figure SI 3). Furthermore, growth of the cultures and overall methylation activity decreased in comparison to BL21, suggesting that BL21*ΔmetJ*-*fcm* was not the optimal host to continue our investigations of *in vivo* methylation from formate. Instead, we turned to strains optimised for formate assimilation based on *E. coli* K-12 MG1655 for introducing formate-derived methyl groups. In the strains C_1_-opt and C_1_S, *fcm* was introduced under control of a constitutive promoter and genes involved in the native generation of C_1_-H_4_F compounds were deleted,^[34]^ leading to excellent result in converting formate into the methyl group of methionine.^[34]^ Complete isolation of the C_1_-H_4_F metabolism was engineered in the C_1_ auxotrophic strain C_1_-opt, where *glyA* and *gcvTHP* were deleted.^[34]^ A partially isolated C_1_-H_4_F metabolism was created in the serine-C_1_ auxotrophic strain C_1_S in which *gcvTHP* was deleted, *glyA* is still intact but the *de novo* biosynthesis of serine is not possible due to knockout of the 3-phospho-glycerate dehydrogenase (*Ec*SerA).^[34]^ An introduced arabinose-inducible T7 RNA-polymerase system alloes overproduction of MTs from the pET28a vector (Figure 2, Table 1).^[44]^

To elucidate basic methylation activity, we focussed on *N*-methylation with *Rg*ANMT and compared BL21-*fcm-rganmt*, C_1_-opt-*rganmt* and C_1_S-*rganmt*. Following the overexpression phase in LB-media, all strains methylate 2,5-ANP to M-2,5-ANP without the addition of formate as analysed with HPLC-UV and LC-MS/MS (Figure 4, Figure SI 1 & SI 4). Product formation with C_1_S-*rganmt* was 1.5-fold higher than with BL21-*rganmt*, while C_1_-opt-*rganmt* exhibited only about 10% methylation activity (Figure 4, Table SI 6). Growth of the cultures in LB-media presumably allowed a certain part of metabolites (*e.g.* methionine, SAM, or serine for C_1_S) to be carried over or reside in the cells enabling *in vivo* methylation activity without the addition of formate. Supplying C_1_-opt-*rganmt* with 5 mM ^13^C-formate led to a 2-fold increase of methylation activity with a share of 54% ^13^C-labelled product. As conversion rates were still only about 5%, this strain was seen as unfit to compete with BL21-*fcm-rganmt* in terms of transformation of formate to methyl groups and overall methylation activity. Methylation activity of C_1_S-*rganmt* increased 1.4-fold (42% total conversion) when 5 mM ^13^C-formate were added, clearly outperforming BL21-*fcm-rganmt-ecmetF* (21% total conversion). To our surprise, only 15% of M-2,5-ANP were ^13^C-labelled, indicating only a fraction of transferred methyl groups to be derived from formate. Due to the high increase in methylation activity, a much higher share was expected. Still, in comparison to BL21-*fcm-rganmt, in vivo* methylation activity with C_1_S-*rganmt* benefitted from the addition of formate which is why we continued focussing on the C_1_-auxotrophic strain.

**Figure 4.**
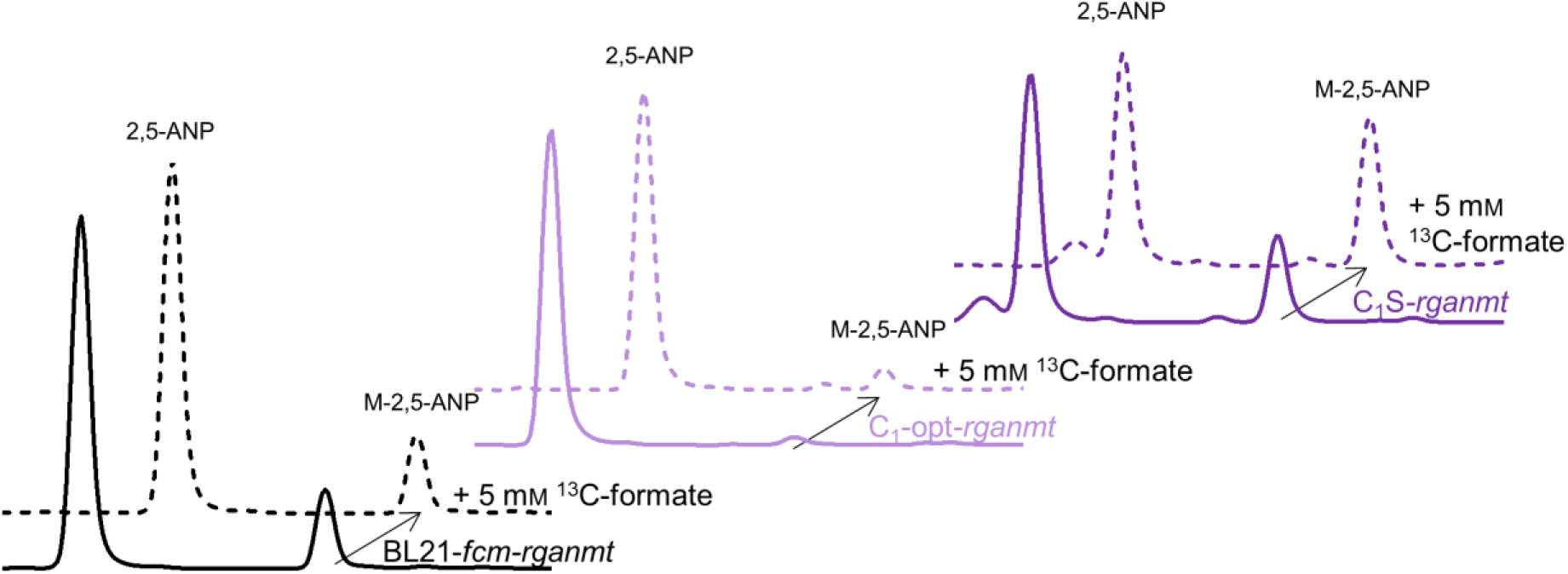
*In vivo* methylation activity of BL21-*fcm-rganmt*, C_1_-opt-*rganmt* and C_1_S-*rganmt*. HPLC-UV (λ = 280 nm) chromatogram of the *in vivo* methylation experiments of BL21-*fcm-rganmt*, C_1_-opt-*rganmt* and C_1_S-*rganmt* with and without the addition of 5 m_M_ ^13^C-formate. Experiments were conducted in M9-medium (22 m_M_ glucose) with a final OD_600_ of 3.0, 0.75 m_M_ 2,5-ANP and 5 m_M_ ^13^C-formate at 37 °C, 170 rpm for 24 h.

To evaluate the influence of formate concentration on *in vivo* methylation activity during the biotransformation phase, formate concentrations ranging from 2.5 mM to 75 mM were screened (Figure 5, Figure SI 4). Substantial changes in conversion started from 2.5 mM and peaked at 10 mM formate supply with a 2.8-fold higher product formation compared to BL21*-fcm-rganmt*.

**Figure 5.**
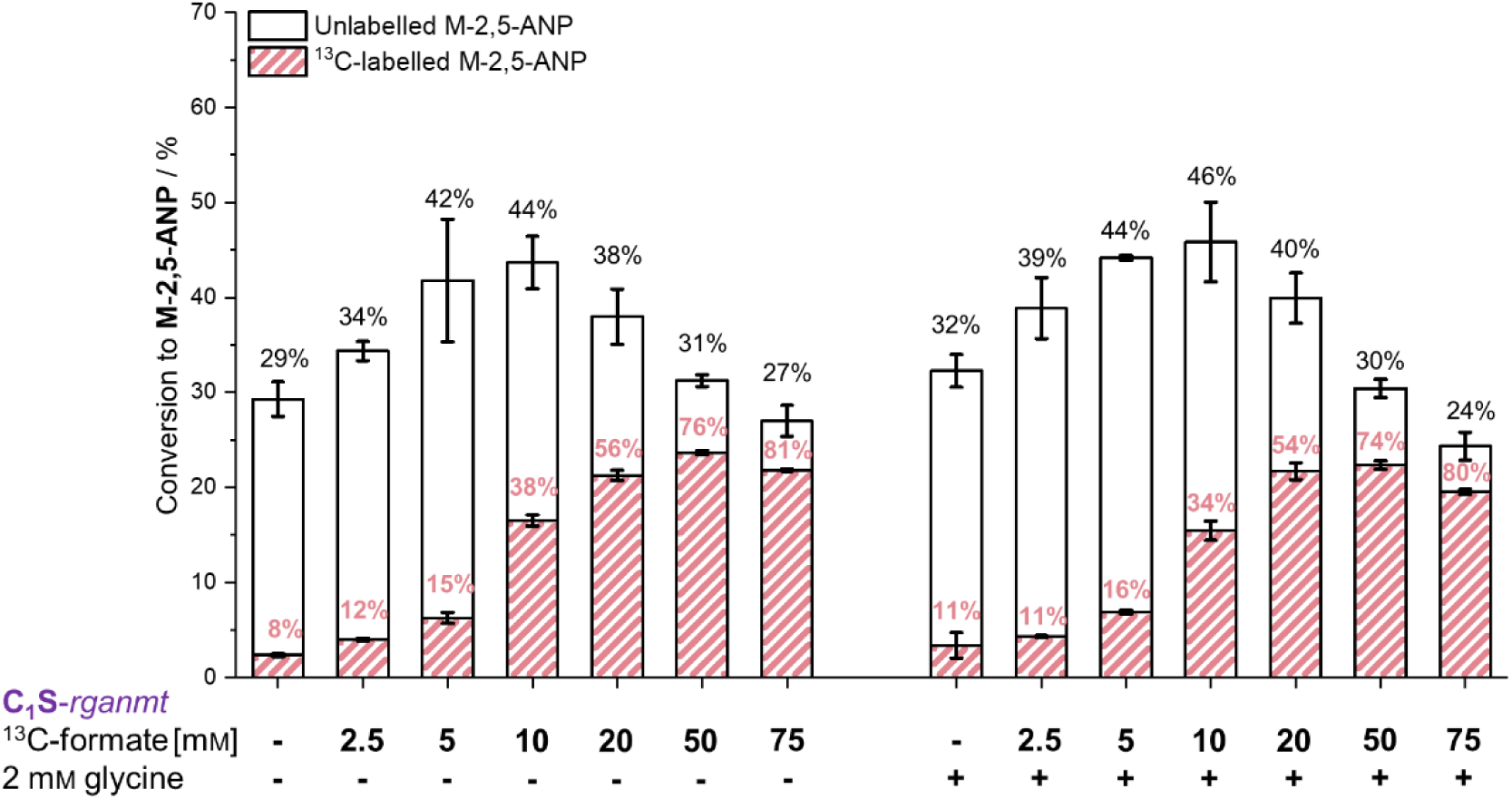
Increasing *in vivo* methylation with formate in C_1_S-rganmt. Given is the conversion of 2,5-ANP to M-2,5-ANP by C_1_S-*rganmt* (black) and the respective share of ^13^C-labelled product (red) after supply with 2.5 – 75 m_M_ ^13^C-formate with and without the addition of 2 m_M_ glycine. Experiments were conducted in M9-medium (22 m_M_ glucose) with a final OD_600_ of 3.0, 0.75 m_M_ 2,5-ANP, 2.5 m_M_ – 75 m_M_ ^13^C-formate and 2 m_M_ glycine at 37 °C, 170 rpm for 24 h. NOTE: About 8% of M--2,5-ANP is naturally ^13^C-labelled, therefore respective shares are indicators not absolutes for formate-derived methyl groups. * = p < 0.05

Shares of labelled product with 10 mM ^13^C-formate did not exceed 38% with 10 mM formate and 2 mM glycine. Further increase of formate concentration lowered methylation activity whereas the share of labelled product increased, indicating a toxic effect of high formate concentrations on C_1_S-*rganmt*. When 50 mM formate were supplied, 31% conversion with 76% ^13^C-labelled methylated product were detected, giving the highest total amount of labelled product.

In the C_1_S strain, *Ec*GlyA is still present and could use methylene-H_4_F to synthesise serine from glycine when an excess of glycine is supplemented. On the one hand, this reaction could withdraw C_1_-compounds from the C_1_-H_4_F metabolism, leading to decreased availability of C_1_-carbon units for *in vivo* methylation activity. On the other hand, allowing serine synthesis could increase cysteine concentrations and through the transsulfurylation pathway also methionine availability, supporting *in vivo* methylation activity.^[45]^ To test this, we repeated the screening with 2.5 mM to 75 mM formate in combination with supply of 2 mM glycine during the biotransformation phase in M9 media (Figure 5). Co-feeding of formate and glycine increased methylation activity slightly (5-10% higher conversion) for every condition up to 20 mM formate. With further increase of the C_1-_compound concentration, benefits of supplied glycine subsided. In most conditions, the share of ^13^C-labelled product was slightly lower when glycine was added, indicating that some of the ^13^C-labelled C_1_-compound was indeed withdrawn from the C_1_-H_4_F metabolism. However, the differences observed were not significant, and supply of 10 mM ^13^C-formate and 2 mM glycine yielded the highest total methylation activity within this study (46% conversion with a share of 34% ^13^C-M-2,5-ANP). Increasing the share of formate-derived methyl groups using low formate concentrations did not prove to be fruitful (Figure SI 6 & SI 7). We therefore focussed on improving total *in vivo* methylation activity of C_1_S-*rganmt* with 50 mM ^13^C-formate.

To do so, the genes of the SAM regeneration cycle as well as *ecmetF* were co-overexpressed on a separate pCDF-DUET-1 vector in C_1_S-*rganmt* (Figure 6). Increasing SAM synthesis by co-overexpression of *ecmat* lowered *in vivo* methylation activity, similar to the experiments with BL21-*rganmt*. Similar results were obtained for the co-overexpression of *ecmetF*. Co-overexpression of *ecmtan* and *ecluxS*, however, led to a slight increase (∼10%) in methylation activity, indicating that removal of SAH and regeneration to homocysteine is one minor factor limiting *in vivo* methylation activity in C_1_S-*rganmt*. Final *in vivo* methylation activity with C_1_S-*rganmt*-*ecmtan*-*ecluxs* supplied with 50 mM ^13^C-formate was more than doubled compared to BL21-*rganmt* and allowed more than 70% of the methyl groups to derive from the renewable C_1_-compound.

**Figure 6.**
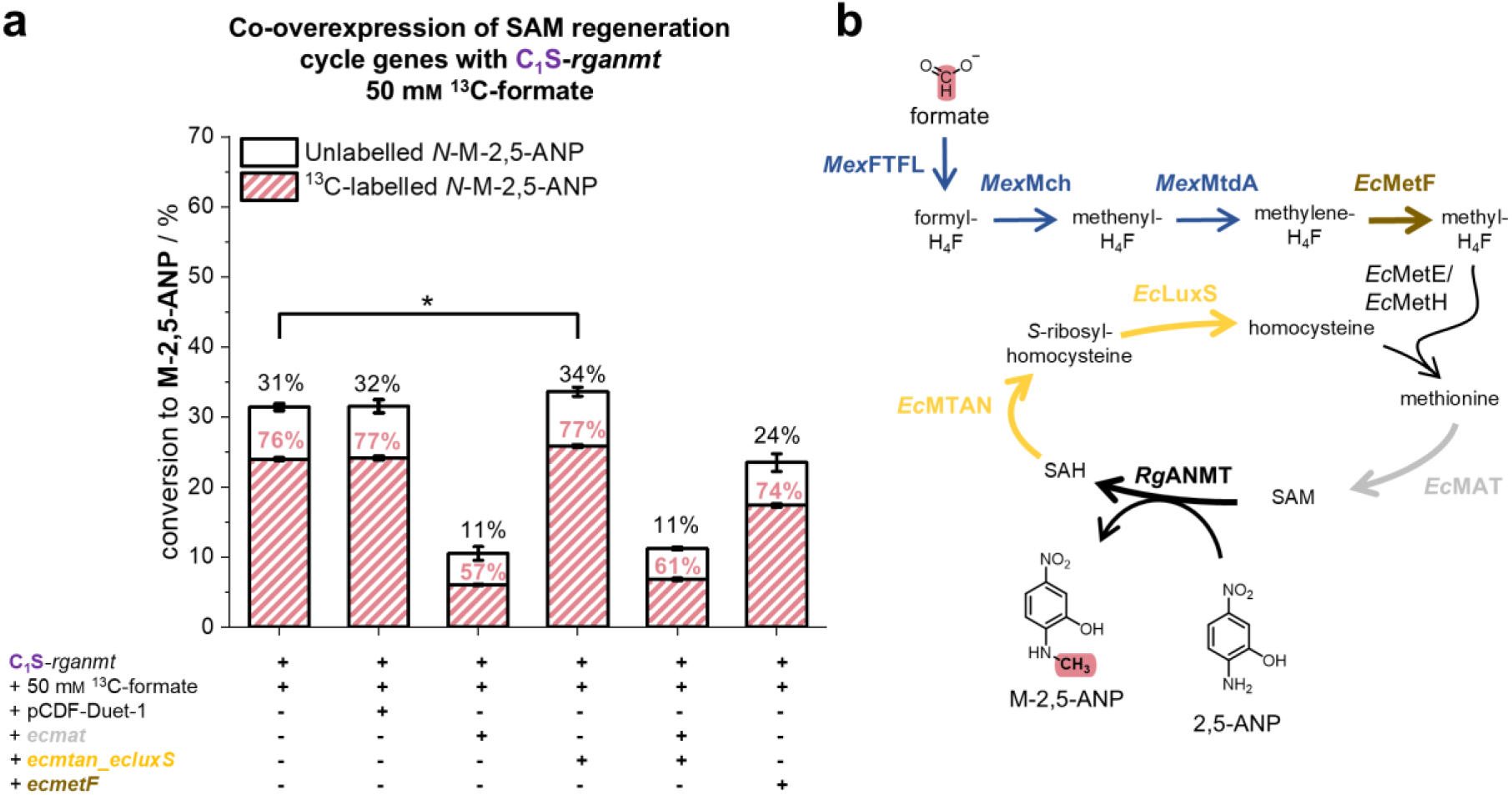
Co-overexpression of SAM regeneration cycle genes in C_1_S-*rganmt*. **a** Co-overexpression of *rganmt* and adjuvant genes encoded on a pCDF-DUET-1 vector in C_1_S. Given are the conversion rates (black) and the relative share of ^13^C-labelled product (red). **b** Scheme of the metabolic pathway from formate to methyl groups in combination with highlighting of the here co-overproduced proteins. Experiments were conducted in M9-medium, with 22 m_M_ glucose, a final OD_600_ of 3.0, 0.75 m_M_ 2,5-ANP and 50 m_M_ ^13^C-formate at 37 °C, 170 rpm for 24 h. NOTE: About 8% of M--2,5-ANP is naturally ^13^C-labelled, therefore respective shares are indicators not absolutes for formate-derived methyl groups. * = p < 0.05

## Conclusion

With these data, we demonstrated introduction of the renewable C_1_-compound formate into SAM-derived methyl groups. Utilisation of pFCM in BL21-*fcm-rganmt* allowed methylation with 67% of ^13^C-labelled product, which was also applicable to the MTs *Pp*CaOMT and CouO. Application of BL21-*fcm*-*mt* offers a simple opportunity to convert formate into methyl groups, allowing for one of the first applications of the feedstock formate as a methylation agent.

We further investigated the *E. coli* strains C_1_-opt and C_1_S with an engineered C_1_-carbon metabolism for straight-forward transformation of formate to methyl groups. Complete isolation of the C_1_-H_4_F metabolism in C_1_-opt resulted in low *in vivo* methylation activity. The serine-dependent C_1_-auxotrophic C_1_S strain on the contrary, exhibited strong methylation activity. Addition of 10 mM formate and 2 mM glycine allowed a 57% increase in methylation with a share of 34% ^13^C-labelled product, allowing formate to not only be used as a methyl group donor but also as an enhancer of *in vivo* methylation activity.

Further increase in formate concentration lowered total *in vivo* methylation activity due to the toxicity of formate, however, increased the share of ^13^C-labelled methylated product. We suspect that this phenomenon is caused by a source of metabolites producing competing C_1_-compounds. One explanation could be given by the synthesis of endogenous formate by the native pyruvate formate-lyase (*Ec*PFL). *Ec*PFL is an oxygen-sensitive enzyme and native expression of *ecpfl* occurs under anaerobic conditions providing a further electron sink for fermentation by cleavage of pyruvate to acetyl-CoA and formate.^[46]^ Another possibility could be the production of intracellular formaldehyde, which can be used by the unspecific threonine aldolase (*Ec*LtaE) to synthesise unlabelled serine, representing a substrate for the production of unlabelled methylene-H_4_F by *Ec*GlyA.^[47]^

Assimilation of formate, reduction to methyl groups and synthesis of SAM is furthermore an energy-consuming process, demanding two equivalents of ATP and two equivalents of NAD(P)H. Synthesis of homocysteine from aspartate further requires three equivalents of ATP and two equivalents of NADH, adding SAM synthesis to the energy intensive formate assimilation.^[21]^ Increasing NADH supply by implementation of, *e.g.* formate dehydrogenase could further improve the intracellular availability of reducing power in *E. coli* and, as a result, also *in vivo* methylation.

Radical SAM enzymes are involved in the synthesis of many pharmaceutically active natural compounds, however, due to their sensitive [4Fe-4S]-cluster, they are difficult to use *in vitro*. Formate-driven regeneration of SAM could also be used to enhance *in vivo* activity of radical SAM MTs and allow harnessing of their potential in the future.

## Author contributions

MKFM and JNA designed the research idea, AS, SLM and TJE further refined the research idea, MKFM performed and analysed *in vivo* methylation experiments, AS prepared the T7-RNA polymerase arabinose-inducible C_1_-opt and C_1_S strains. MKFM wrote and together with JNA revised the manuscript, all authors contributed together to elaboration of the final manuscript.

## Acknowledgements

We thank Aaliya Afandi Lee and Adelheid Nagel for their skilful technical support. We thank Prof. Dr Nicholas Turner for providing the *E. coli* BL21*ΔmetJ* strain. We thank Dr Hai He for providing progenitor strains used in the construction of C_1_-opt and C_1_S. This work was funded by a PhD scholarship of the Deutsche Bundesstiftung Umwelt (DBU, grant number 20021/742) to MKFM and the European Research Council (ERC, grant number 716966). JNA acknowledges funding through a Heisenberg Grant of the Deutsche Forschungsgemeinschaft (DFG/527572100).

## Materials and methods

### Material

All chemicals were purchased in a high purity from Sigma Aldrich, Carl Roth or Thermo Scientific, except stated otherwise. ^13^C-formate (>99% ^13^C-enriched) was obtained from Sigma Aldrich. Unlabelled formate was obtained from Carl Roth chemicals.

### Strains and plasmids used in this study

**Table 1.**
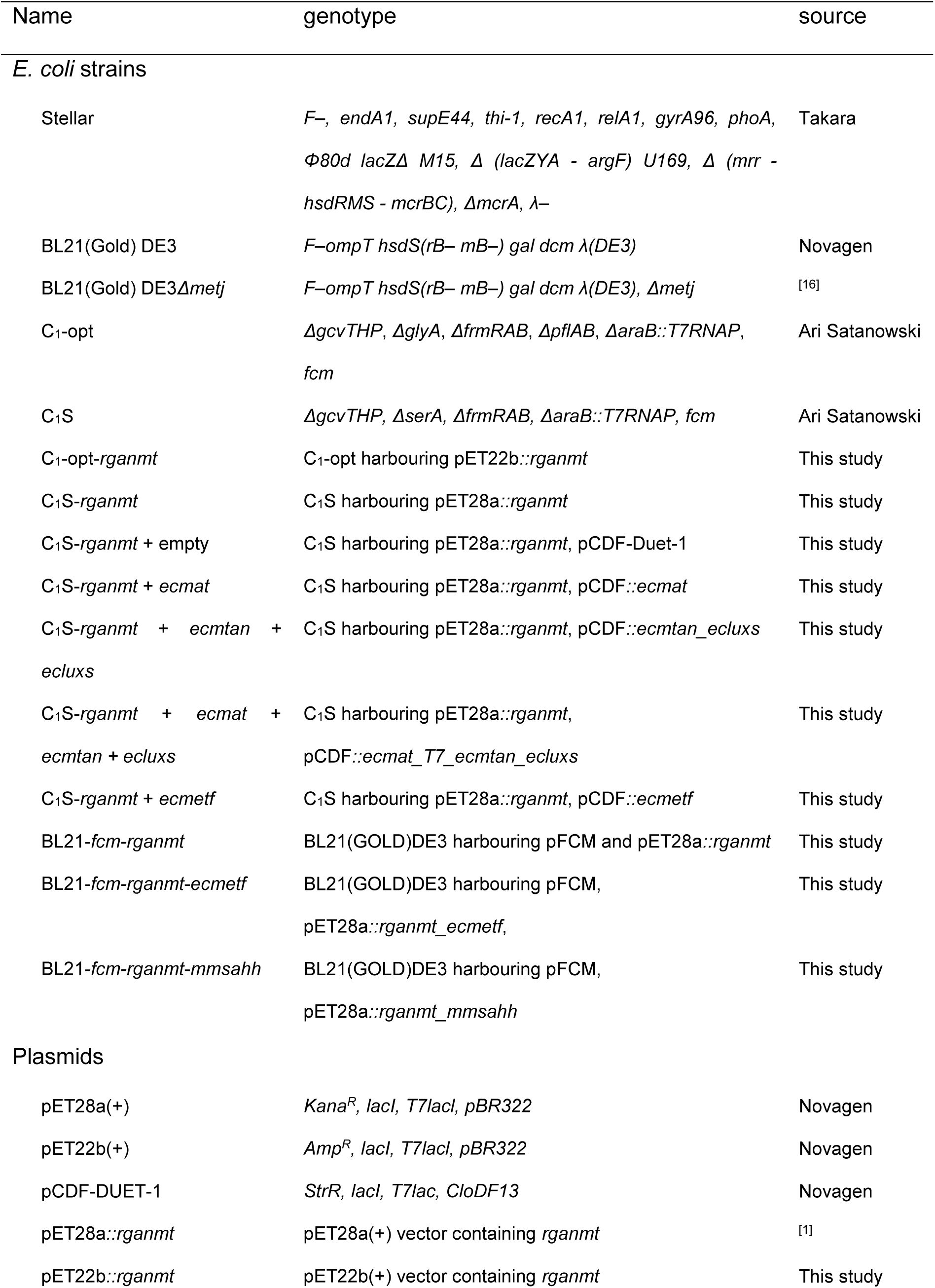
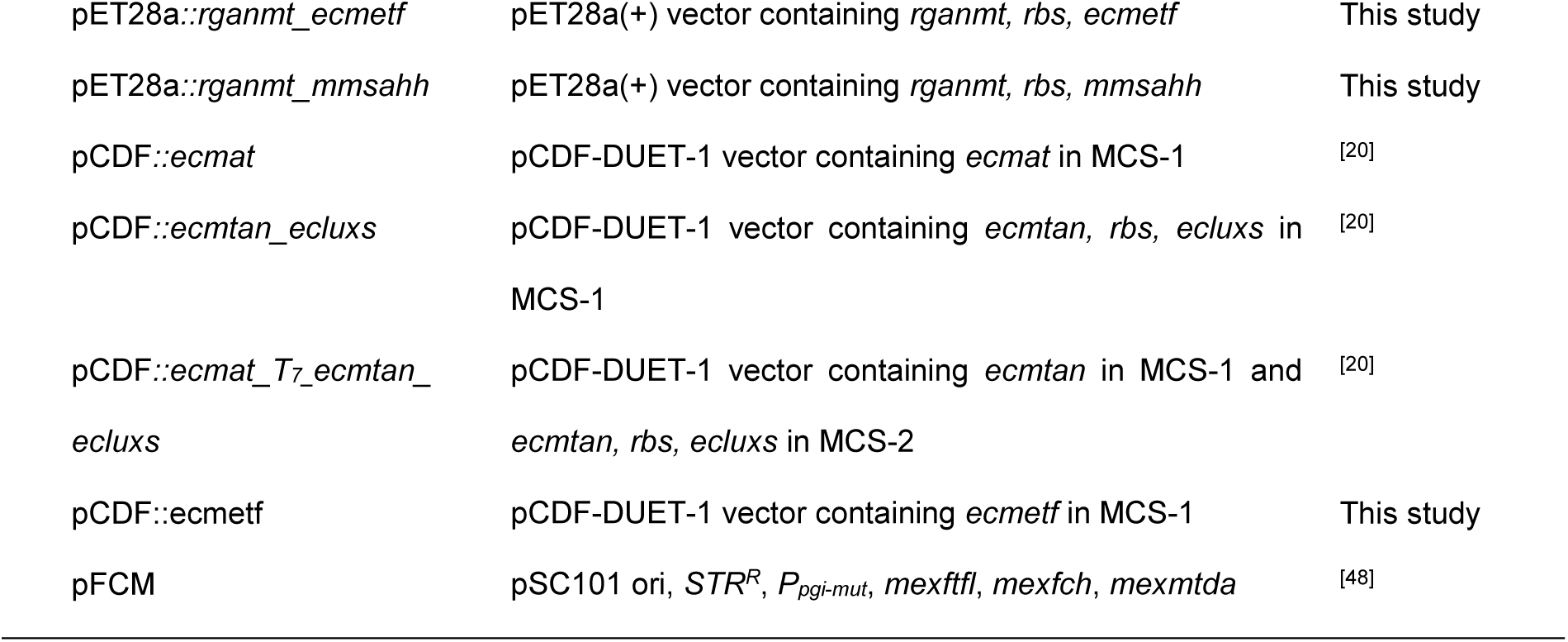
Strains and plasmids used in this study.

## Methods

### *E. coli* strain engineering

All strains used in this study are listed in Table 1. The strains C_1_-opt and C_1_S were constructed from *E. coli* strain SIJ488,^[49]^ a derivative of MG1655, engineered to carry inducible genes for λ-Red recombineering and flippase in its chromosome. Genome engineering was performed by λ-red recombineering^[50]^ and P1 phage transduction.^[49,50]^ In brief, the genes indicated for each strain in Table 1 were deleted using PCR-amplified linear DNA fragments carrying kanamycin or chloramphenicol resistance markers flanked by FRT-sites based on the pKD4 or pKD3 plasmids, respectively.^[50]^ Resistance markers were amplified with primers carrying 50bp overhangs serving as homologous regions flanking the target gene.^[51]^ For removal of resistance markers from the chromosome, flippase was induced by addition of L-rhamnose as described previously.^[49]^ Gene deletions and removal of resistance cassettes were verified by PCR.

For chromosomal insertion of the genes encoding the enzymes for formate assimilation (*Mex*FTFL, *Mex*Fch and *Mex*MtdA, together referred to as FCM),^[9,28]^ we used a previously described genome insertion protocol^[52]^ and vector containing the desired genes (“pDM4:SS9-C_1_M”).^[9]^ In brief, a non-replicative plasmid (pDM4, R6K ori) was introduced into the respective recipient strain via conjugation from an *E. coli* ST18 donor strain. Kanamycin resistance was used to select for chromosomal insertion of the synthetic FCM operon (including the kanamycin resistance gene) via native homologous recombination based on 600bp homology regions, with subsequent levansucrase (*sacB*) counter-selection.

For chromosomal insertion of the arabinose-inducible T7-RNA polymerase, we transferred the corresponding genomic locus from BL21-AI (Invitrogen) using P1 phage transduction. The transferred locus consists of an insertion replacing the araB gene, introducing genes encoding a tetracycline efflux pump (a selectable tetracycline resistance marker) and T7-RNA polymerase.^[44]^The latter is controlled by the endogenous P_araBAD_ promoter, thus achieving arabinose-inducible expression of T7-RNA polymerase. Successful transductants were selected via tetracycline resistance and the insertion site was verified by PCR.

### Cloning procedure

C_1_-opt already contains a Kana^R^ resistance, which is why *rganmt* was cloned with the restriction enzymes *NdeI* and *HindIII* into an *Amp^R^*containing pET22b(+) vector, linearised with the same restriction enzymes.

For co-overexpression on pET28a vector with BL21, *ecmetf* and *mmsahh* were cloned behind *rganmt* by digesting pET28a*::rganmt* with *XhoI* and cloning of the respective insert into the linearised vector. When co-overexpression was conducted on pET28 and pCDF vector in C_1_S, *ecmetf* was cloned into the first MCS of pCDF-DUET-1. For this purpose, the respective inserts were amplified from a pET28a vector *via* polymerase chain reaction (Table 2) and cloned into the linearised vectors by in-fusion cloning.

Primers with 15 bp overhangs complementary to the vector were ordered at Eurofins as lyophilised powder. The PCR was set up with 3 µl fwrd primer (c = 10 pM), 3 µl rev primer (c = 10 pM, Table 2), 1 µl of template (pET28a vector with the gene of interest or yeast gDNA), 18 µl nuclease free water, and 25 µl 2x Phusion MasterMix (Thermo Fisher). The PCR was conducted with 5 min initial denaturation at 98 °C, followed by 29 cycles of 20 s denaturation at 98 °C, 20 s annealing at 55 °C and 35 s elongation at 72°C. Afterwards, 5 min terminal elongation at 72 °C followed. The size of the gene was confirmed by agarose gel electrophoresis (1% agarose, 100 V, 50 min), followed by extraction of the band and clean up with the Promega Wizard™ Plus DNA Purification System. For insertion of the gene, the respective vectors were linearised with the respective primers or with *XhoI* (Table 2). 50 ng of the linear vector were combined with a 1:3 ratio of the gene to be inserted, 2 µl of In-Fusion 5x buffer in a total volume of 10 µl nuclease free water. The mixture was incubated at 50 °C for 20 min followed by standard chemical transformation into *E. coli* Stellar cells. Identity of each vector was confirmed by sequencing.

**Table 2.**
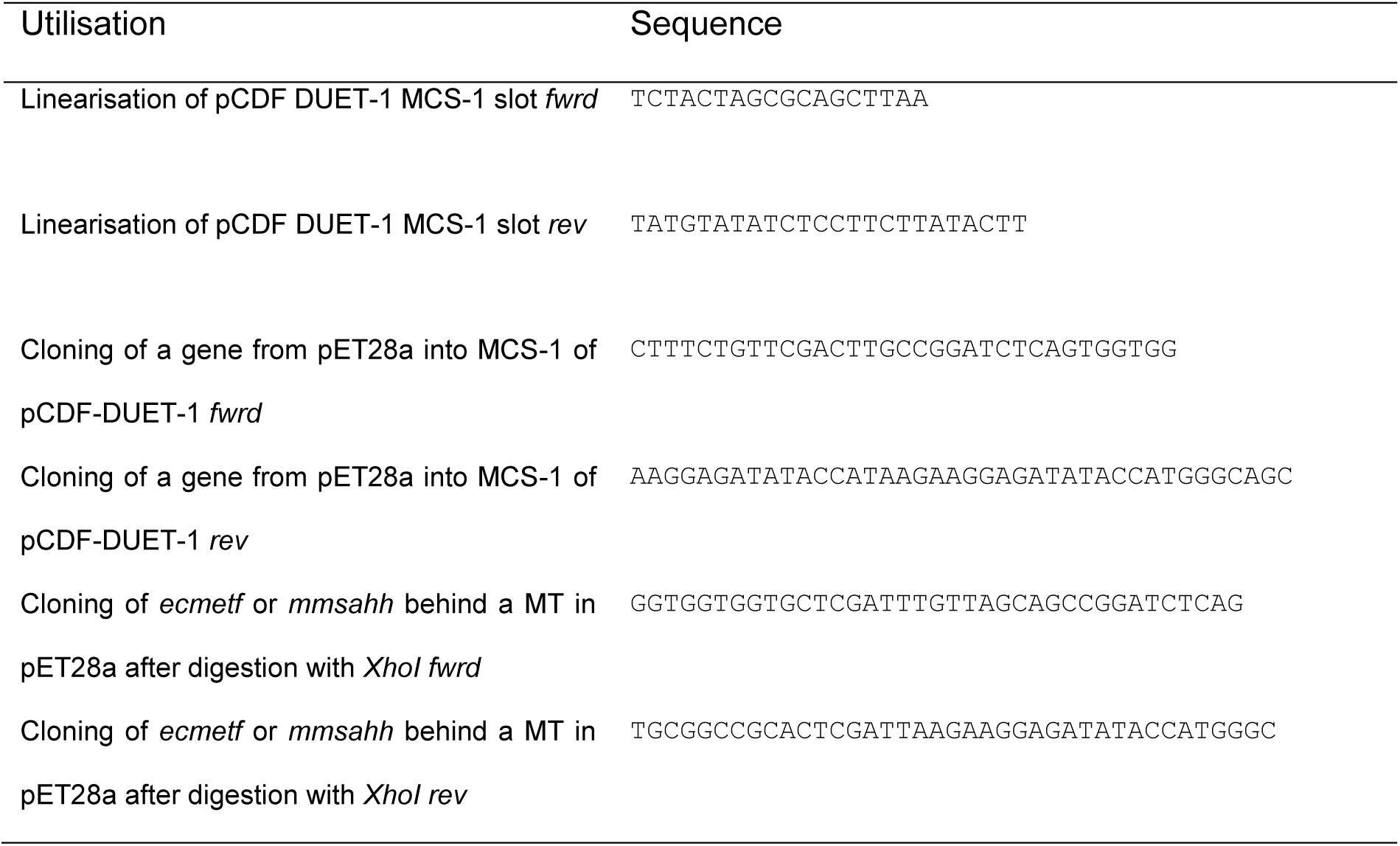
Utilised oligonucleotides for vector cloning.

### *In vivo* assay procedure

For *in vivo* experiments, the respective plasmids were chemically (co)-transformed in the respective *E. coli* strain and grown overnight at 37 °C on agar plates containing the respective antibiotics. A single colony was picked and used to inoculate 5 ml LB-medium containing the respective antibiotic followed by overnight incubation at 37 °C, 170 rpm. The preculture was used to inoculate 25 ml LB-medium containing the respective antibiotic in a 50 ml falcon tube in a ratio of 1:100. The culture was grown to an OD_600_ of 0.5 – 0.7, overexpression was fully induced with 1 mM IPTG and took place at 20 °C, 140 rpm for 20 h.

After overexpression, the cultures were chilled on ice for 10 min followed by centrifugation for 15 min, 4 °C, 2,500 g. The pellet was washed with 5 ml of ice cold M9-medium (0.4% glucose, 20 mM glycerol or 13 mM xylose, 48 mM Na_2_HPO_4_, 22 mM KH_2_PO_4_, 8.6 mM NaCl, 1.8 mM NH_4_Cl, 2 mM MgSO_4_, 0.1 mM CaCl_2_) followed by centrifugation 15 min, 4 °C, 2,500 g.

The pellet was then resuspended to an OD_600_ of approximately 4.5 in ice cold M9-medium containing 1 mM IPTG and the respective antibiotics. Assays were then pipetted on ice in 1.5 ml reaction tubes each consisting of 1 ml of the assay mixture followed by incubation at 37 °C and 170 rpm. The whole cell catalysts were used in a final OD_600_ of 3.0 with 0.75 mM of 2,5-ANP, 1.25 mM – 50 mM (^13^C-)formate and 2 mM glycine. Incubation took place for 24 h at 37 °C and 170 rpm. For the analysis of methylation activity,100 µl of the assay mixture was stopped after 24 h with 35 µl of 10% perchloric acid followed by centrifugation 4 °C and 18,000 g for 30 min. The methylated product was quantified *via* HPLC-UV (λ = 280 nm) and the share of ^13^C-labelled product was elucidated by LC-MS/MS.

### Statistics

To test for significances a two-sided t-test of two independent samples of the same variance was used. A significant difference in activity was accepted with p < 0.05. All experiments were conducted at least in biological triplicates.

### HPLC-UV analysis

HPLC-UV analysis was used to detect MT products generated during *in vivo* methylation.

**Table 3.** HPLC-UV analysis for quantification of methylated products.

|  | HPLC-UV analysis for quantification of M-ANP |
| --- | --- |
| HPLC system | Agilent 1260 infinity II |
| Column | ISAspher 100-3-C18 (150 x 4.0 mm) |
| Mobile phase A (aqueous) | 0.1% formic acid |
| Mobile phase B (organic) | acetonitrile |
| Detection wavelength | 280 nm |
| Flow rate | 1 ml/ min |
| Sample volume injected | 10 µl |
| Equilibration | 97% A |
| Gradient | 4 min 97% A/ 3% B |
|  | 3 min to 30% A/ 70% B |
|  | 0.1 min to 100% B |
|  | 2.9 min 100% B |
|  | 1 min to 97% A/ 3% B |
|  | 3 min 97% A/ 3% B |

### LC-MS/MS analysis

LC-MS/MS was used for analysis of the share of ^13^C-labelled products.

**Table 4.** LC-MS/MS method for determination of ^13^C-labelled products.

| | LC-MS/MS method for determination of $^{13}\text{C}$-labelled M-ANP |
| --- | --- |
| LC-MS system | SCIEX 5500+ |
| Column | ISAspher 120-3-C18 Aq (50 x 2.1 mm) |
| Mobile phase A (aqueous) | 0.1% formic acid |
| Mobile phase B (organic) | acetonitril |
| Flow rate | 0.2 ml/ min |
| MS settings | Curtain Gas: 25 psi<br>Ion spray voltage: +2900 eV<br>Temperature: 450 °C<br>Ion source gas 1: 50 psi<br>Ion source gas 2: 50 psi<br>Declustering potential: 10 eV<br>Exit potential: 12 eV |
| Sample preparation | Dilution 1:10 and filtration |
| Sample volume injected | 1 µl |
| Equilibration | 97% A |
| Gradient | 4 min 97% A/ 3% B<br>3 min to 30% A/ 70% B<br>0.1 min to 100% B<br>2.9 min 100% B<br>1 min to 97% A/ 3% B<br>3 min 97% A/ 3% B |

**Table 5.** Multiple reaction monitoring transitions for LC-MS/MS measurements.

| Substrate | Q1 mass | Q3 mass |
| --- | --- | --- |
| ANP | 155.050 | 155.050 |
|  | 155.050 | 109.050 |
| M-ANP | 169.050 | 169.050 |
|  | 169.050 | 123.050 |
| $^{13}\text{C}$-M-ANP | 170.050 | 170.050 |
|  | 170.050 | 124.050 |
| 2,4-MNA | 169.050 | 169.050 |
|  | 169.050 | 123.050 |
| $^{13}\text{C}$-2,4-MNA | 170.050 | 170.050 |
|  | 170.050 | 124.050 |
| DHN | 161.050 | 161.050 |
| Methyl-DHN | 175.070 | 175.070 |
| $^{13}\text{C}$-Methyl-DHN | 176.070 | 176.070 |

## References

[1] Y. Kloo, L. J. Nilsson, E. Palm, Renew. Sustain. Energy Transit. 2024, 5, 100075.

[2] L. Wollensack, K. Budzinski, J. Backmann, Curr. Opin. Green Sustain. Chem. 2022, 33, 100586.

[3] A. Bazzanella, D. Krämer, DECHEMA e.V. 2017.

[4] R. A. Sheldon, J. M. Woodley, Chem. Rev. 2018, 118, 801–838.

[5] T. Schaub, Phys. Sci. Rev. 2018, 3, 3.

[6] P. Germer, J. N. Andexer, M. Müller, Synthesis 2022, 54, 4401–4425.

[7] O. Yishai, S. N. Lindner, J. Gonzalez De La Cruz, H. Tenenboim, A. Bar-Even, Curr. Opin. Chem. Biol. 2016, 35, 1–9.

[8] M. Nattermann, S. Wenk, P. Pfister, H. He, S. H. Lee, W. Szymanski, N. Guntermann, F. Zhu, L. Nickel, C. Wallner, J. Zarzycki, N. Paczia, N. Gaißert, G. Franciò, W. Leitner, R. Gonzalez, T. J. Erb, Nat. Commun. 2023, 14, 2682.

[9] S. Kim, S. N. Lindner, S. Aslan, O. Yishai, S. Wenk, K. Schann, A. Bar-Even, Nat. Chem. Biol. 2020, 16, 538–545.

[10] A.-W. Struck, M. L. Thompson, L. S. Wong, J. Micklefield, ChemBioChem 2012, 13, 2642–2655.

[11] S. Mordhorst, J. N. Andexer, Nat. Prod. Rep. 2020, 37, 1316–1333.

[12] E. Abdelraheem, B. Thair, R. F. Varela, E. Jockmann, D. Popadić, H. C. Hailes, J. M. Ward, A. M. Iribarren, E. S. Lewkowicz, J. N. Andexer, P. Hagedoorn, U. Hanefeld, ChemBioChem 2022, e202200212.

[13] H. Schönherr, T. Cernak, Angew. Chem. Int. Ed. 2013, 52, 12256–12267.

[14] C. R. Kirman, L. M. Sweeney, M. L. Gargas, J. H. Kinzell, Inhal. Toxicol. 2009, 21, 537– 551.

[15] J. B. Broderick, W. E. Broderick, B. M. Hoffman, FEBS Lett. 2023, 597, 92–101.

[16] J. L. Galman, F. Parmeggiani, L. Seibt, W. R. Birmingham, N. J. Turner, Angew. Chem. Int. Ed. 2022, 61, e202112855.

[17] Q. Liu, B. Lin, Y. Tao, Metab. Eng. 2022, 72, 46–55.

[18] B. David, J.-L. Wolfender, D. A. Dias, Phytochem. Rev. 2015, 14, 299–315.

[19] Z. W. Luo, J. S. Cho, S. Y. Lee, Proc. Natl. Acad. Sci. 2019, 116, 10749–10756

[20] M. Mohr, P. Bencic, J. N. Andexer, Angew. Chem. Int. Ed. 2024, e202414598.

[21] Y. Lv, J. Chang, W. Zhang, H. Dong, S. Chen, X. Wang, A. Zhao, S. Zhang, Md. A. Alam, S. Wang, C. Du, J. Xu, W. Wang, P. Xu, J. Agric. Food Chem. 2024, 72, 3846– 3871.

[22] K. Okano, Y. Sato, S. Inoue, S. Kawakami, S. Kitani, K. Honda, Catalysts 2020, 11.

[23] J. C. Gonzalez, R. V. Banerjee, S. Huang, J. S. Sumner, R. G. Matthews, Biochem. 1992, 31, 6045–6056.

[24] P. Simic, J. Willuhn, H. Sahm, L. Eggeling, Appl. Environ. Microbiol. 2002, 68, 3321– 3327.

[25] G. Kikuchi, Mol. Cell. Biochem. 1973, 1, 169–187.

[26] J. W. Locasale, Nat. Rev. Cancer 2013, 13, 572–583.

[27] T. Sahr, S. Ravanel, F. Rébeillé, Biochem. Soc. Trans. 2005, 33, 758–762.

[28] S. Wenk, V. Rainaldi, K. Schann, H. He, M. Bouzon, V. Döring, S. N. Lindner, A. Bar-Even, Metab. Eng. 2024, DOI 10.1016/j.ymben.2024.10.007.

[29] S. Kim, S. H. Lee, H. Seo, K.-J. Kim, Biochem. Biophys. Res. Commun. 2020, 528, 426–431.

[30] A. M. Ochsner, F. Sonntag, M. Buchhaupt, J. Schrader, J. A. Vorholt, Appl. Microbiol. Biotechnol. 2015, 99, 517–534.

[31] J. A. Vorholt, L. Chistoserdova, M. E. Lidstrom, R. K. Thauer, J. Bacteriol. 1998, 180, 5351–5356.

[32] S. Yilmaz, B. Kanis, R. A. H. Hogers, S. Benito-Vaquerizo, J. Kahnt, T. Glatter, B. Dronsella, T. J. Erb, M. Suarez-Diez, N. J. Claassens, bioRxiv 2024, 2024.11.11.622960.

[33] C. A. Sheppard, E. E. Trimmer, R. G. Matthews, J. Bacteriol. 1999, 181, 718–725.

[34] O. Yishai, L. Goldbach, H. Tenenboim, S. N. Lindner, A. Bar-Even, ACS Synth. Biol. 2017, 6, 1722–1731.

[35] B. Berger, M. Knodel, BMC Microbiol 2003, 3, 12.

[36] N. Parveen, K. A. Cornell, Mol. Microbiol. 2011, 79, 7–20.

[37] K. Winzer, K. R. Hardie, N. Burgess, N. Doherty, D. Kirke, M. T. G. Holden, R. Linforth, K. A. Cornell, A. J. Taylor, P. J. Hill, P. Williams, Microbiology 2002, 148, 909–922.

[38] S. Shimizu, S. Shiozaki, T. Ohshiro, H. Yamada, Eur. J. Biochem. 1984, 141, 385–392.

[39] E. Jockmann, F. Subrizi, M. K. F. Mohr, E. M. Carter, P. M. Hebecker, D. Popadić, H. C. Hailes, J. N. Andexer, ChemCatChem 2023, 15, e202300930.

[40] B. Lin, Y. Tao, Microb. Cell Factories 2017, 16, 106.

[41] C. C. C. R. de Carvalho, Microb. Biotechnol. 2017, 10, 250–263.

[42] F. Marincs, I. W. Manfield, J. A. Stead, K. J. Mcdowall, P. G. Stockley, Biochem. J. 2006, 396, 227–234.

[43] R. Shoeman, T. Coleman, B. Redfield, R. C. Greene, A. A. Smith, I. Saint-Girons, N. Brot, H. Weissbach, Biochem. Biophys. Res. Commun. 1985, 133, 731–739.

[44] N. Bhawsinghka, K. F. Glenn, R. M. Schaaper, Microbiol. Resour. Announc. 2020, 9, 10.1128/mra.00009-20.

[45] S. M. Aitken, P. H. Lodha, D. J. K. Morneau, Biochim. Biophys. Acta BBA - Proteins Proteomics 2011, 1814, 1511–1517.

[46] G. Sawers, A. Böck, J. Bacteriol. 1989, 171, 2485–2498.

[47] H. He, R. Höper, M. Dodenhöft, P. Marlière, A. Bar-Even, Metab. Eng. 2020, 60, 1–13.

[48] O. Yishai, M. Bouzon, V. Döring, A. Bar-Even, ACS Synth. Biol. 2018, 7, 2023–2028.

[49] S. I. Jensen, R. M. Lennen, M. J. Herrgård, A. T. Nielsen, Sci. Rep. 2015, 5, 17874.

[50] K. A. Datsenko, B. L. Wanner, Proc. Natl. Acad. Sci. 2000, 97, 6640–6645.

[51] T. Baba, T. Ara, M. Hasegawa, Y. Takai, Y. Okumura, M. Baba, K. A. Datsenko, M. Tomita, B. L. Wanner, H. Mori, Mol. Syst. Biol. 2006, 2, 2006.0008.

[52] S. Wenk, O. Yishai, S. N. Lindner, A. Bar-Even, in Methods Enzymol. (Ed.: N. Scrutton), Academic Press, 2018, pp. 329–367.

