## Supplemental Information for "Rewiring *Escherichia coli* to Transform Formate into Methyl Groups"

### Supplementary figures & data

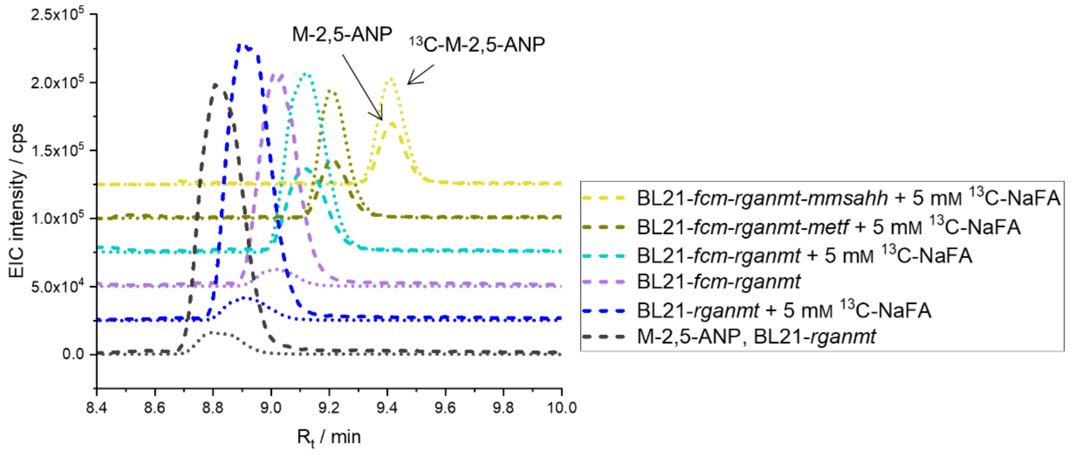


**Figure SI 1. Analysis of formate derived methyl groups with BL21-*fcm-rganmt*.** Shown is the LC-MS/MS analysis of $^{13}$C-labelled product with BL21-*fcm-rganmt* and strains co-overexpressing genes from the SAM regeneration cycle or *ecmetf*, when 5 mM $^{13}$C-formate was supplied. Experiments were conducted in M9-medium (22 mM glucose) with a final $OD_{600}$ of 3.0, 0.75 mM of substrate to be methylated, 0 - 5 mM $^{13}$C-formate at 37 °C, 170 rpm for 24 h. Experiments were conducted in biological triplicates.

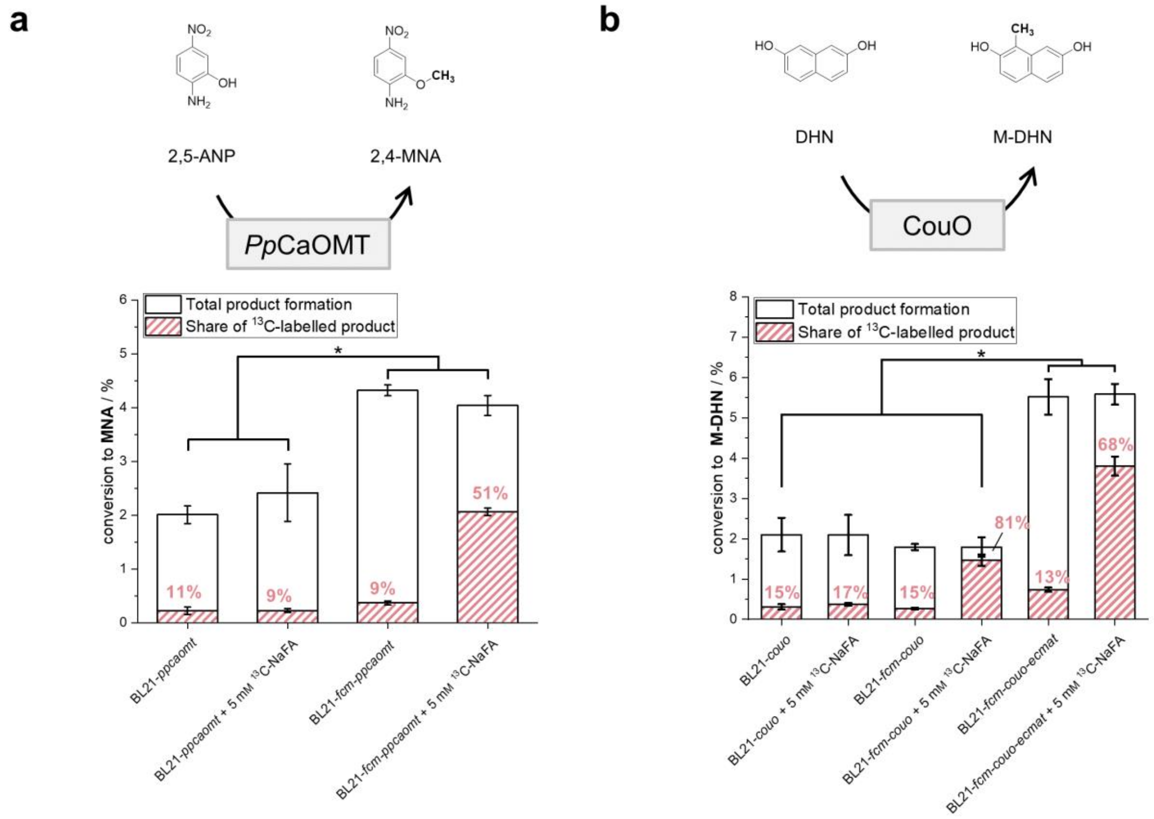


**Figure SI 2. General applicability of methylation from formate with BL21.** Bar charts show production of methylated products with BL21-*ppcaomt* (**a**) or BL21-*couo* (**b**) in combination with *fcm* and supply with 5 mM $^{13}$C-formate. Experiments were conducted in M9-medium (22 mM glucose) with a final $OD_{600}$ of 3.0, 0.75 mM of substrate to be methylated, 0 - 5 mM $^{13}$C-formate at 37 °C, 170 rpm for 24 h. Experiments were conducted in biological triplicates.

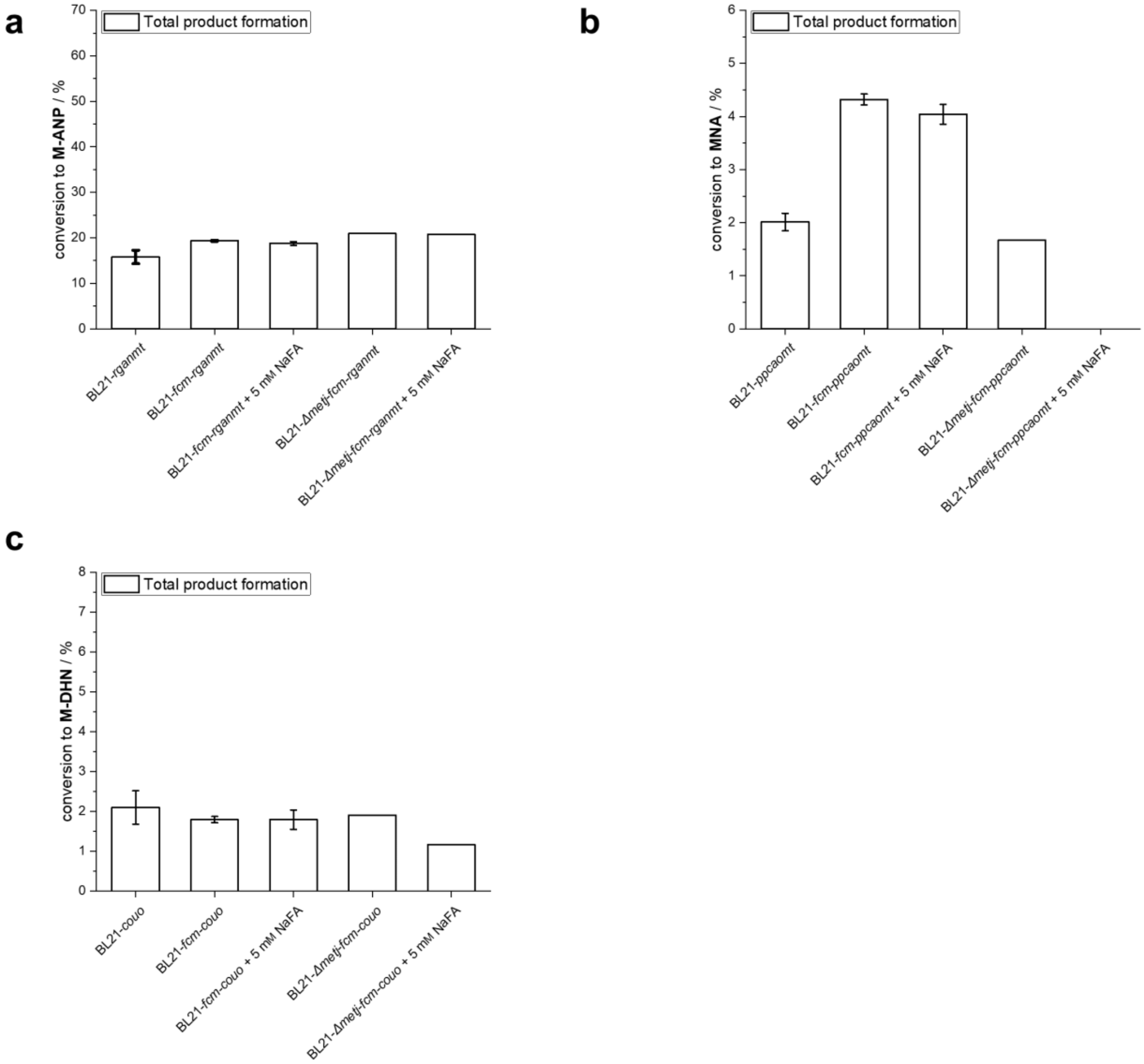


**Figure SI 3. Comparison of *in vivo* methylation from formate with BL21*::Δmetj-fcm* and BL21-*fcm*.** Shown is product formation of the strains bearing **a** *Rg*ANMT, **b** *Pp*CaOMT and **c** CouO in BL21, BL21-*fcm* and BL21*::Δmetj-fcm* with and without the addition of formate. Experiments were conducted in M9-medium (22 mM glucose) with a final $OD_{600}$ of 3.0, 0.75 mM of substrate to be methylated, 0 - 5 mM $^{13}$C-formate at 37 °C, 170 rpm for 24 h. NOTE: in this experiment no $^{13}$C-labelled formate could be used. Experiments were performed in biological triplicates.

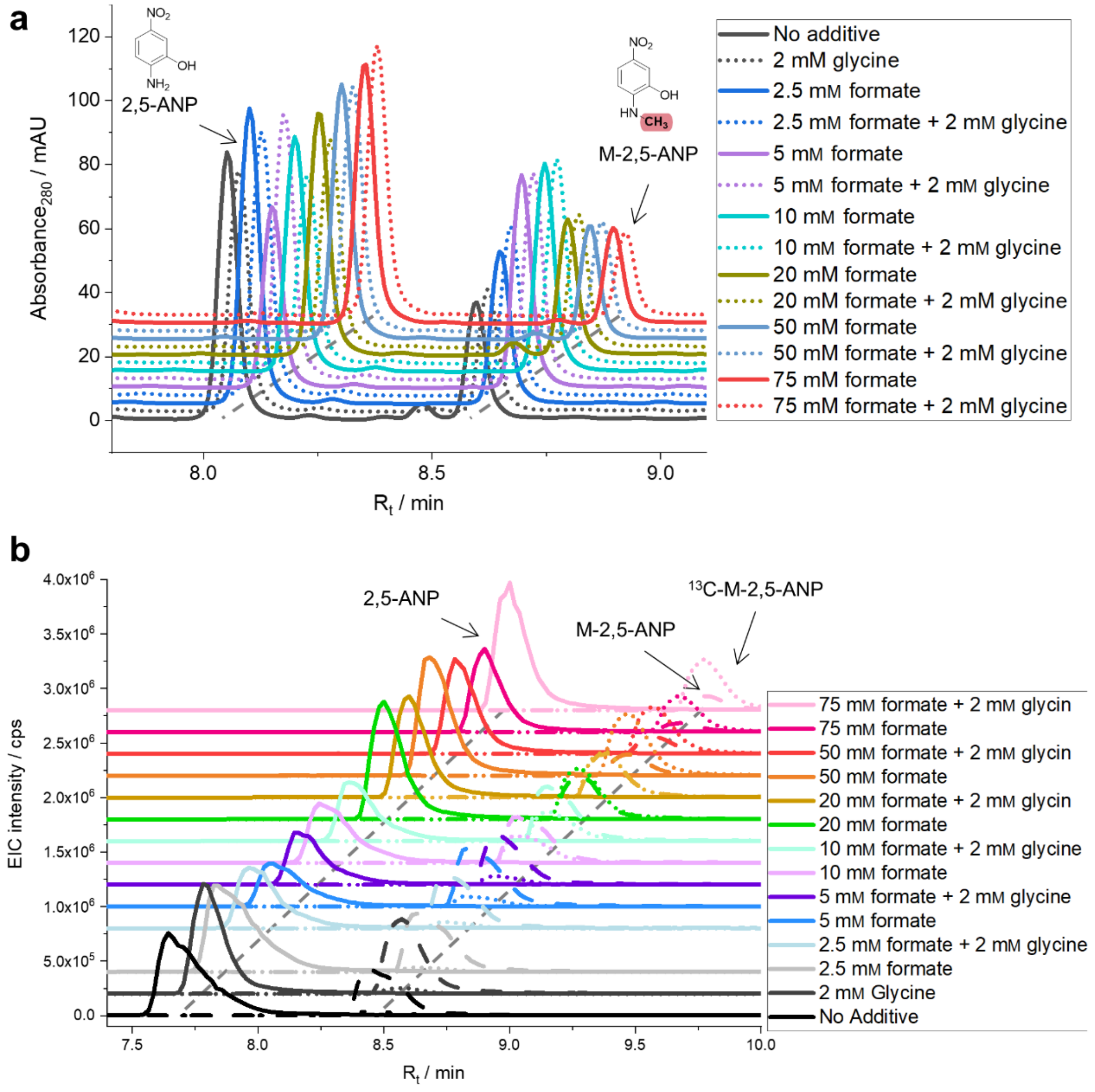


**Figure SI 4. Screening of different $^{13}$C-formate concentrations with $C_1$S-*rganmt*. a** Production of M-2,5-ANP was analysed with HPLC-UV (λ = 280 nm) using $C_1$S-*rganmt* with formate concentrations ranging from 2. 5 to 75 mM $^{13}$C-formate with and without the addition of 2 mM glycine. **b** LC-MS/MS analysis of the incorporation of formate derived methyl groups in M-2,5-ANP. Experiments were conducted in M9-medium (22 mM glucose) with a final $OD_{600}$ of 3.0, 0.75 mM ANP, 2.5 - 75 mM $^{13}$C-formate with or without 2 mM glycine at 37 °C, 170 rpm for 24 h in biological triplicates.

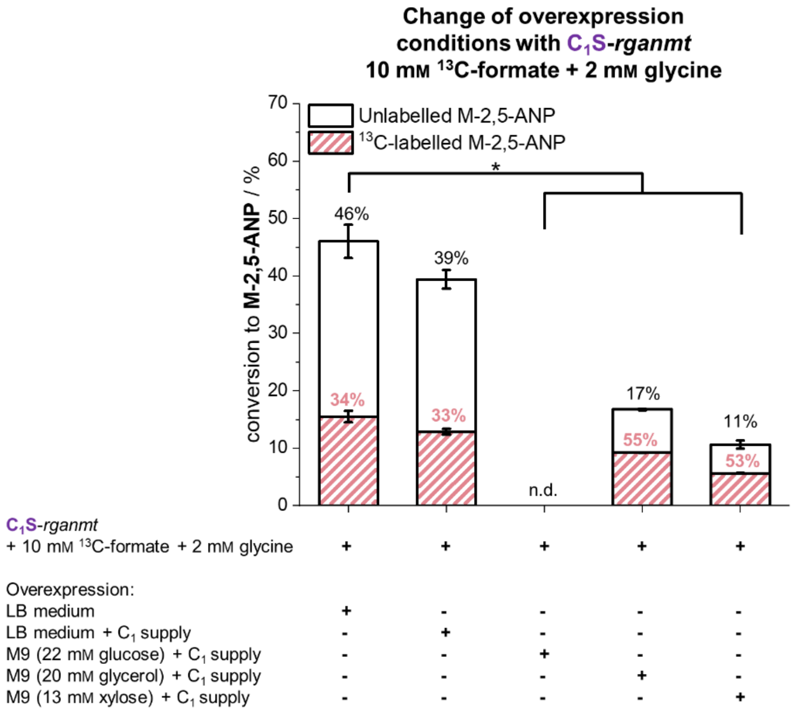


**Figure SI 5. Exchange of overexpression conditions for $C_1$S-*rganmt*.** Given is the conversion of 2,5-ANP to M-2,5-ANP by $C_1$S-*rganmt* (black) and the share of $^{13}$C-labelled product (red). Conditions during overexpression and biotransformation phase were altered as indicated to M9 (22 mM glucose, 20 mM glycerol, or 13 mM xylose) at a final $OD_{600}$ of 3.0, 0.75 mM 2,5-ANP, 10 mM $^{13}$C-formate and 2 mM glycine at 37 °C, 170 rpm for 24 h.

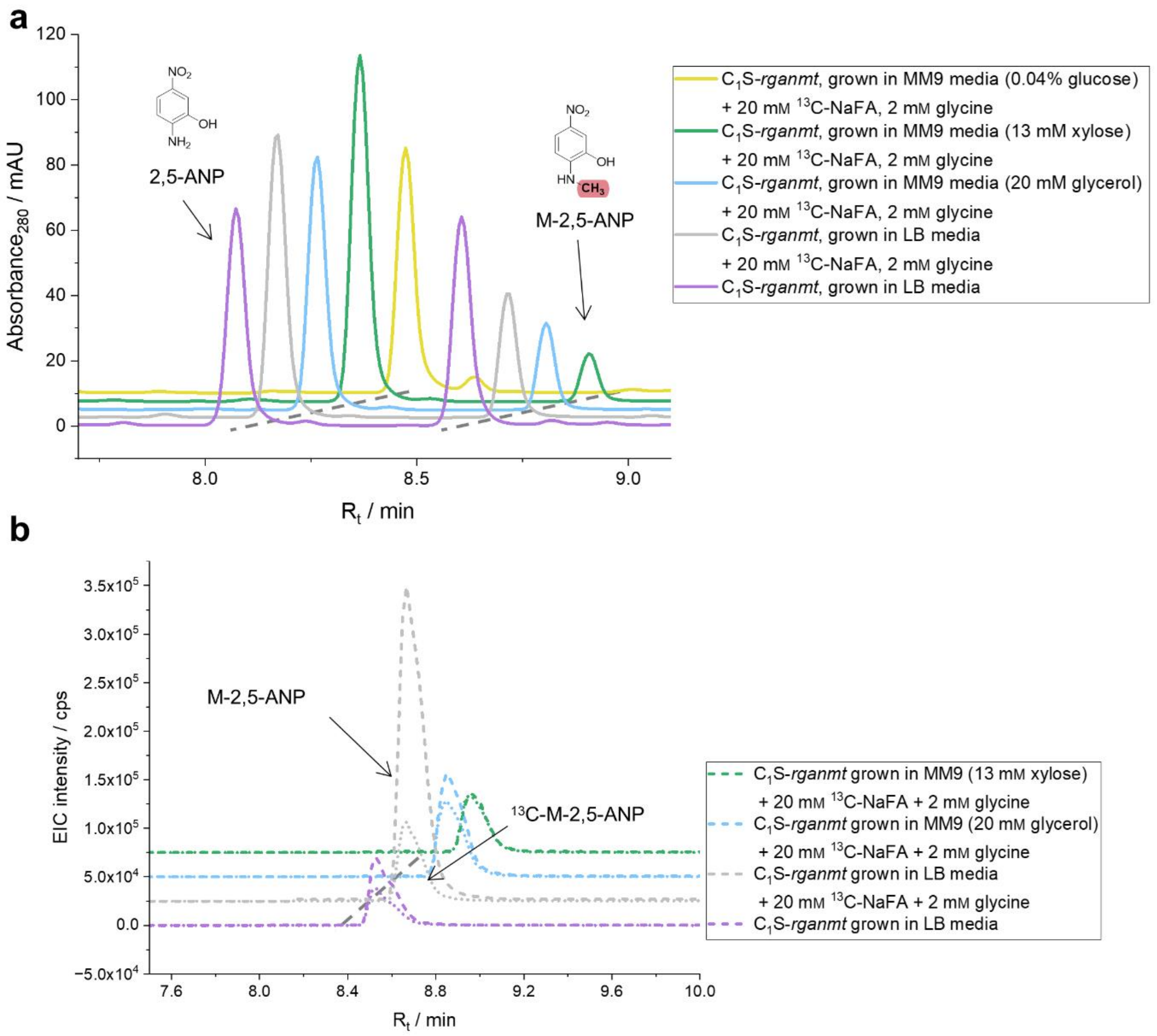


**Figure SI 6. Media screening to increase methylation from formate with $C_1$S-*rganmt*. a** Total methylation of 2,5-ANP to M-2,5-ANP through $C_1$S-*rganmt* grown and overexpressed in different medium was analysed with HPLC-UV (λ = 280 nm). **b** The share of labelled product was analysed *via* LC-MS/MS. Striped lines represent M-2,5-ANP (m/z = 169.050-123.050) and dotted lines represent $^{13}$C-M-2,5-ANP (m/z = 170.050-124.050). Cells were grown and overexpressed in the respective medium and the experiments were conducted in M9-medium (22 mM glucose or with the respective M9-medium the cultures were grown in) with a final $OD_{600}$ of 3.0, 0.75 mM 2,5-ANP, 10 mM $^{13}$C-formate with 2 mM glycine at 37 °C, 170 rpm for 24 h in biological triplicates.

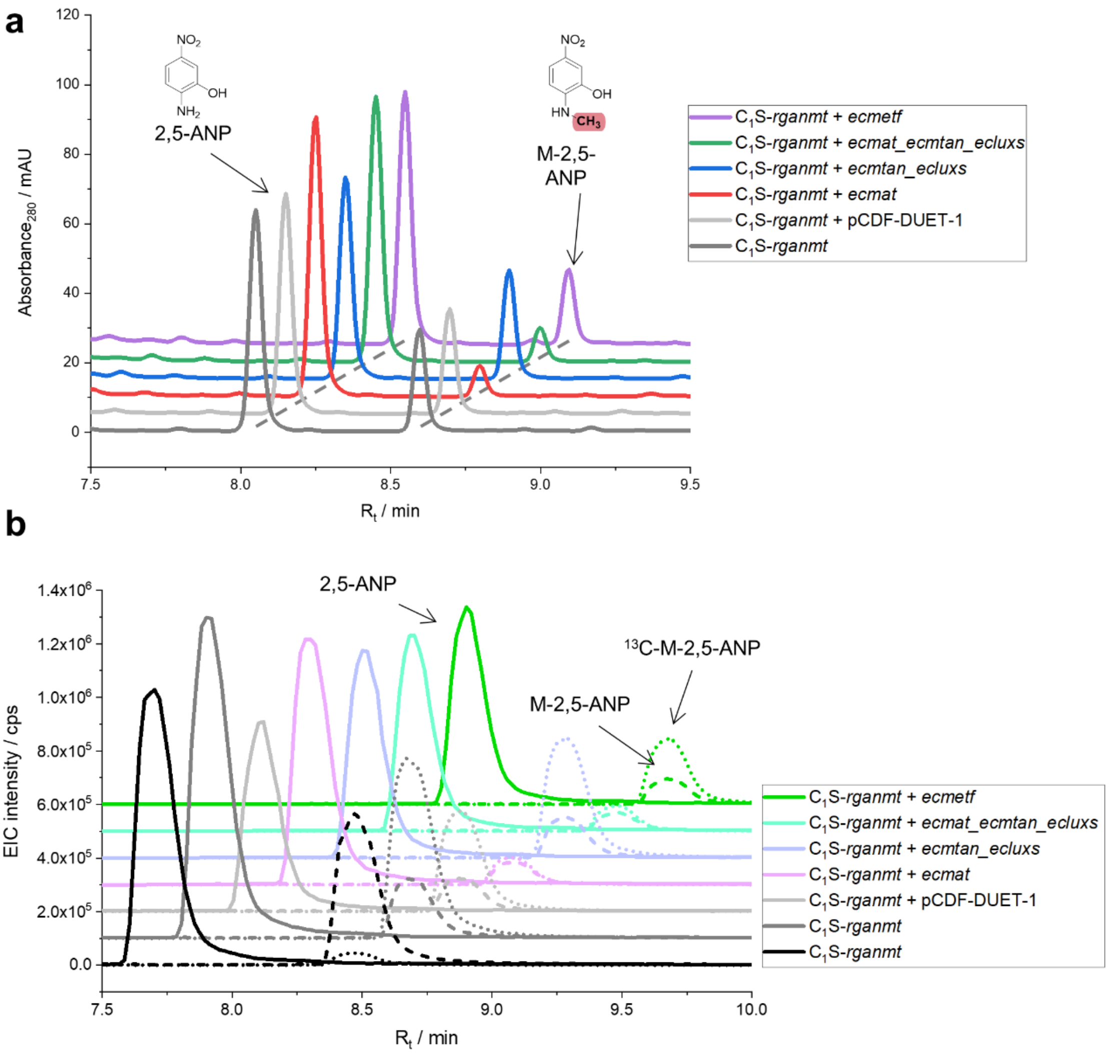


**Figure SI 7. Co-overexpression of SAM regeneration cycle genes in $C_1$S-*rganmt*. a** Production of M-2,5-ANP was analysed with HPLC-UV (λ = 280 nm) with $C_1$S-*rganmt* in combination with the SAM regeneration cycle genes encoded on a pCDF--DUET-1 vector using 50 mM $^{13}$C-formate. **b** Incorporation of $^{13}$C-formate was analysed with LC-MS/MS. Compact lines represents 2,5-ANP (m/z = 155.050-109.050), striped lines represent M-2,5-ANP (m/z = 169.050-123.050) and dotted lines represent $^{13}$C-M-2,5-ANP (m/z = 170.050-124.050). Experiments were conducted in M9-medium (22 mM glucose) with a final $OD_{600}$ of 3.0, 0.75 mM 2,5-ANP, and 50 mM $^{13}$C-formate at 37 °C, 170 rpm for 24 h in biological triplicates.

**Table SI 1. Production of M-2,5-ANP for different strains and conditions.**

| Strain | Condition | Conversion ± SD [%] | Fold change ± SD | Share of $^{13}C$-labelled M-2,5-ANP [%] |
|---|---|---|---|---|
| BL21-*rganmt* | - | 15.8 ± 1.5 | 1 ± 0.09 | 7.7 ± 0.4 |
| | 5 mM $^{13}C$-formate | 15.5 ± 0.2 | 0.98 ± 0.01 | 7.8 ± 0.2 |
| BL21-*fcm-rganmt* | - | 19.3 ± 0.1 | 1.22 ± 0.01 | 7.6 ± 0.3 |
| | 5 mM $^{13}C$-formate | 18.7 ± 0.4 | 1.19 ± 0.03 | 67.3 ± 1.4 |
| BL21-*fcm-rganmt-ecmat* | 5 mM $^{13}C$-formate | - | - | - |
| BL21-*fcm-rganmt-mmsahh* | 5 mM $^{13}C$-formate | 17.4 ± 1.3 | 1.10 ± 0.09 | 64.1 ± 1.5 |
| BL21-*fcm-rganmt-ecmeF* | 5 mM $^{13}C$-formate | 20.5 ± 0.8 | 1.30 ± 0.05 | 66.6 ± 1.8 |
| $C_1$-opt-*rganmt* | - | 2.2 ± 0.15 | 0.14 ± 0.01 | 9.1 ± 1.2 |
| | 5 mM $^{13}C$-formate | 5.1 ± 0.15 | 0.32 ± 0.01 | 53.6 ± 0.96 |
| $C_1$S-*rganmt* | - | 29.3 ± 1.8 | 1.9 ± 0.1 | 8.0 ± 0.3 |
| | 2.5 mM formate | 34.4 ± 1.0 | 2.2 ± 0.1 | 11.7 ± 0.4 |
| | 5 mM formate | 41.8 ± 6.4 | 2.6 ± 0.4 | 15.0 ± 1.4 |
| | 10 mM formate | 43.7 ± 2.8 | 2.8 ± 0.2 | 37.8 ± 1.4 |
| | 20 mM formate | 38.0 ± 2.9 | 2.4 ± 0.2 | 56.0 ± 1.4 |
| | 50 mM formate | 31.2 ± 0.6 | 2.0 ± 0.1 | 75.7 ± 0.7 |
| | 75 mM formate | 27.0 ± 1.6 | 1.7 ± 0.1 | 80.8 ± 0.4 |
| | 2 mM glycine | 32.3 ± 1.7 | 2.0 ± 0.1 | 10.5 ± 4.1 |
| | 2.5 mM formate + 2 mM glycine | 38.9 ± 3.2 | 2.5 ± 0.2 | 11.1 ± 0.2 |
| | 5 mM formate + 2 mM glycine | 44.2 ± 0.3 | 2.8 ± 0.1 | 15.5 ± 0.4 |
| | 10 mM formate + 2 mM glycine | 46.0 ± 4.2 | 2.9 ± 0.3 | 33.7 ± 2.2 |
| | 20 mM formate + 2 mM glycine | 40.0 ± 2.6 | 2.5 ± 0.2 | 54.3 ± 2.2 |
| | 50 mM formate + 2 mM glycine | 30.4 ± 1.0 | 1.9 ± 0.1 | 73.5 ± 1.5 |
| | 75 mM formate + 2 mM glycine | 24.3 ± 1.5 | 1.5 ± 0.1 | 80.3 ± 1.15 |
| $C_1$S-*rganmt* | Overexpression in LB-medium + 20 mM $^{13}C$-formate + 2 mM glycine, assay in M9 (22 mM glucose); 10 mM $^{13}C$-formate + 2 mM glycine | 39.4 ± 1.6 | 2.49 ± 0.10 | 32.7 ± 1.28 |
| $C_1$S-*rganmt* | Overexpression in M9 (22 mM glucose) + 20 mM $^{13}C$-formate + 2 mM glycine, assay in M9 (22 mM glucose) 10 mM $^{13}C$-formate + 2 mM glycine | - | - | - |
| $C_1$S-*rganmt* | Overexpression in M9 (20 mM glycerol) + 20 mM $^{13}C$-formate + 2 mM glycine, assay in M9 (20 mM glycerol) 10 mM $^{13}C$-formate + 2 mM glycine | 16.8 | 1.06 ± 0.01 | 55.0 ± 0.18 |

| | | | | |
|---|---|---|---|---|
| $C_1$S-*rganmt* | Overexpression in M9 (13 mM xylose) + 20 mM $^{13}$C-formate + 2 mM glycine, assay in M9 (13 mM xylose) 10 mM $^{13}$C-formate + 2 mM glycine | 10.6 ± 0.7 | 0.67 ± 0.04 | 53.2 ± 0.6 |
| $C_1$S-*rganmt-empty* | 50 mM $^{13}$C-formate | 31.5 ± 0.9 | 2.0 ± 0.1 | 76.6 ± 0.8 |
| $C_1$S-*rganmt-ecmat* | 50 mM $^{13}$C-formate | 10.5 ± 1.0 | 0.7 ± 0.1 | 57.3 ± 1.1 |
| $C_1$S-*rganmt-ecmtan-ecluxs* | 50 mM $^{13}$C-formate | 33.6 ± 0.6 | 2.1 ± 0.1 | 76.9 ± 0.6 |
| $C_1$S-*rganmt-ecmat-ecmtan-ecluxs* | 50 mM $^{13}$C-formate | 11.3 ± 0.2 | 0.7 ± 0.1 | 60.6 ± 1.6 |
| $C_1$S-*rganmt-ecmetF* | 50 mM $^{13}$C-formate | 23.5 ± 1.3 | 1.5 ± 0.1 | 74.0 ± 1.1 |

### Gene sequences used in this study:

*mexftfl*

ATGCCGAGCGATATTGAAATTGCCCGTGCAGCAACCCTGAAACCGATTGCACAGGTTGCAGAAAAACTGGGTATTCCGGATGAAGCAC
TGCATAACTATGGTAAACATATCGCCAAAATCGACCACGATTTCATTGCAAGCCTGGAAGGTAAACCGGAAGGCAAACTGGTTCTGGT
TACCGCAATTAGCCCGACACCGGCAGGCGAAGGTAAAACCACCACCACCGTTGGTCTGGGTGATGCACTGAATCGTATTGGTAAACGT
GCAGTTATGTGTCTGCGTGAACCGAGCCTGGGTCCGTGTTTTGGTATGAAAGGTGGTGCAGCAGGCGGTGGTAAAGCCCAGGTTGTGC
CGATGGAACAAATCAATCTGCATTTTACCGGTGATTTTCATGCAATTACCAGCGCACATAGCCTGGCAGCAGCACTGATTGATAATCA
TATCTATTGGGCCAACGAGCTGAATATTGATGTTCGTCGTATTCATTGGCGTCGTGTTGTTGATATGAATGATCGTGCACTGCGTGCA
ATTAATCAGAGCTTAGGTGGTGTTGCAAATGGTTTTCCGCGTGAAGATGGTTTTGATATCACCGTTGCAAGCGAAGTTATGGCAGTGT
TTTGTCTGGCCAAAAATCTGGCAGATCTGGAAGAACGTCTGGGTCGTATTGTTATTGCAGAAACCCGTGATCGTAAACCGGTTACACT
GGCAGATGTTAAAGCAACCGGTGCCATGACCGTTCTGCTGAAAGATGCACTGCAGCCGAATCTGGTTCAGACCCTGGAAGGCAATCCG
GCACTGATTCATGGTGGTCCGTTTGCAAATATTGCCCATGGTTGTAATAGCGTTATTGCAACCCGTACCGGTCTGCGTCTGGCCGATT
ATACCGTTACCGAAGCAGGTTTTGGTGCCGATCTGGGTGCCGAAAAATTCATTGACATTAAATGTCGTCAGACGGGTCTGAAACCGTC
AGCAGTTGTTATTGTTGCAACCATTCGTGCCCTGAAAATGCATGGTGGTGTTAATAAGAAAGATCTGCAGGCAGAAAATCTGGATGCA
CTGGAAAAAGGTTTTGCAAATCTGGAACGCCATGTGAATAATGTTCGTAGCTTTGGTCTGCCGGTTGTTGTTGGTGTGAATCATTTCT
TTCAGGATACCGATGCAGAACATGCACGTCTGAAAGAACTGTGTCGTGATCGTCTGCAGGTTGAAGCAATTACCTGTAAACATTGGGC
TGAAGGCGGTGCCGGTGCAGAAGCACTGGCACAGGCCGTTGTTAAACTGGCCGAAGGTGAACAGAAACCGCTGACCTTTGCATATGAA
ACCGAAACCAAAATTACCGACAAAATCAAAGCCATTGCCACCAAACTGTATGGTGCAGCCGATATTCAGATTGAAAGCAAAGCAGCAA
CCAAACTGGCAGGCTTTGAAAAAGATGGTTATGGCAAACTGCCGGTTTGTATGGCAAAAACCCAGTATAGCTTTAGCACCGATCCGAC
ACTGATGGGTGCACCGAGCGGTCATCTGGTTAGCGTTCGTGATGTGCGTCTGAGCGCAGGCGCTGGTTTTGTTGTTGTTATTTGTGGT
GAAATCATGACCATGCCAGGTCTGCCGAAAGTTCCGGCAGCAGATACCATTCGTCTGGATGCCAATGGTCAGATTGATGGTCTGTTTT
AA

*mexfch*

ATGGCCGGCAACGAGACGATCGAAACATTCCTCGATGGCCTGGCGAGCTCGGCCCCGACCCCCGGCGGCGGCGGTGCCGCCGCGATCT
CCGGCGCCATGGGCGCGGCGCTGGTCTCGATGGTGTGTAACCTCACCATCGGCAAGAAGAAGTATGTCGAGGTCGAGGCCGACCTGAA
GCAGGTGCTGGAGAAGTCGGAAGGCCTGCGCCGCACGCTCACCGGCATGATCGCCGACGACGTCGAGGCTTTCGACGCGGTGATGGGC
GCCTACGGGCTGCCGAAAAACACCGACGAGGAGAAGGCTGCCCGCGCCGCCAAGATTCAGGAGGCGCTCAAGACCGCGACCGACGTGC
CGCTCGCCTGCTGCCGCGTCTGCCGCGAGGTGATCGATCTGGCCGAGATCGTCGCCGAGAAGGGCAATCTCAACGTCATCTCGGATGC
CGGCGTCGCCGTGCTCTCGGCCTATGCCGGTCTGCGCTCGGCGGCCCTCAACGTCTACGTCAACGCCAAGGGCCTCGACGACCGCGCC
TTCGCCGAGGAGCGGCTGAAGGAGCTGGAAGGCCTGCTGGCCGAGGCGGGCGCGCTCAACGAGCGGATCTACGAGACGGTGAAGTCCA
AGGTAAACTGA

*mexmtda*

ATGTCCAAGAAGCTGCTCTTCCAGTTCGACACCGATGCCACGCCGAGCGTCTTCGACGTCGTCGTCGGCTACGACGGCGGTGCCGACC
ACATCACCGGCTACGGCAACGTCACGCCCGACAACGTCGGCGCCTATGTCGACGGCACGATCTACACCCGCGGCGGCAAGGAGAAGCA
GTCGACGGCGATCTTCGTCGGCGGCGGCGACATGGCGGCCGGCGAGCGGGTGTTCGAGGCGGTGAAGAAGCGCTTCTTCGGCCCGTTC
CGCGTGTCCTGCATGCTGGATTCGAACGGCTCCAACACGACCGCCGCGGCGGGTGTGGCGCTCGTCGTCAAGGCGGCGGGCGGCTCGG
TCAAGGGCAAGAAGGCCGTCGTGCTCGCGGGCACCGGCCCGGTCGGCATGCGCTCGGCGGCGCTGCTCGCCGGCGAGGGCGCCGAGGT
CGTGCTGTGCGGGCGCAAGCTCGACAAGGCGCAGGCCGCGGCCGATTCCGTGAACAAGCGCTTCAAGGTGAACGTCACCGCGGCCGAA
ACCGCGGACGACGCTTCGCGCGCCGAGGCCGTGAAGGGCGCCCATTTCGTCTTCACCGCCGGTGCGATCGGCCTTGAACTGCTGCCGC
AGGCAGCCTGGCAGAACGAGAGTTCGATCGAGATCGTGGCCGACTACAACGCCCAGCCGCCGCTCGGCATCGGCGGGATCGATGCGAC
CGACAAAGGCAAGGAATACGGCGGAAAGCGCGCCTTCGGTGCGCTCGGCATCGGCGGCTTGAAGCTCAAGCTGCACCGCGCCTGCATC
GCCAAGCTGTTCGAGTCGAGCGAAGGCGTCTTCGACGCCGAGGAGATCTACAAGCTGGCCAAGGAAATGGCCTGA

*rganmt* (His$_6$-tagged)

ATGGGCAGCAGCCATCATCATCATCATCACAGCAGCGGCCTGGTGCCGCGCGGCAGCCATATGGGTAGCCTGAGCGAAAGCCATACCC
AGTATAAACATGGTGTTGAAGTGGAAGAAGATGAAGAGGAATCATATAGCCGTGCAATGCAGCTGAGCATGGCAATTGTTCTGCCGAT
GGCAACCCAGAGCGCAATTCAGCTGGGTGTTTTTGAAATTATTGCAAAAGCACCGGGTGGTCGTCTGAGCGCAAGCGAAATTGCAACC
ATTCTGCAGGCACAGAATCCGAAAGCTCCGGTTATGCTGGATCGTATGCTGCGTCTGCTGGTTAGCCATCGTGTTCTGGATTGTAGCG
TTAGCGGTCCGGCAGGCGAACGTCTGTATGGTCTGACCAGCGTTAGCAAATATTTCGTTCCGGATCAGGATGGTGCAAGCCTGGGCAA
TTTTATGGCACTGCCGCTGGATAAAGTTTTTATGGAAAGCTGGATGGGTGTTAAAGGTGCAGTTATGGAAGGTGGTATTCCGTTTAAT
CGTGTTCATGGCATGCACATCTTTGAATATGCAAGCAGCAATAGCAAATTCAGCGATACCTATCATCGTGCCATGTTTAATCATAGCA
CCATTGCACTGAAACGCATCCTGGAACACTATAAAGGTTTTGAGAATGTGACCAAACTGGTTGATGTTGGTGGTGGTCTGGGTGTTAC
CCTGAGCATGATTGCAAGCAAATATCCGCATATTCAGGCCATCAATTTTGATCTGCCGCATGTTGTTCAGGATGCAGCAAGCTATCCG
GGTGTTGAACATGTGGGTGGTAATATGTTTGAAAGCGTTCCGGAAGGTGATGCCATTCTGATGAAATGGATTCTGCATTGTTGGGATG
ATGAACAGTGTCTGCGTATTCTGAAAAACTGCTATAAAGCCACACCGGAAAATGGCAAAGTGATTGTTATGAATAGCGTTGTTCCGGA
AACACCGGAAGTTAGCAGCAGCGCACGTGAAACCAGCCTGCTGGATGTTCTGCTGATGACCCGTGATGGTGGTGGCCGTGAACGTACC
CAGAAAGAATTTACCGAACTGGCAATTGGTGCAGGTTTTAAAGGTATTAACTTTGCCTGTTGCGTGTGCAACCTGCATATTATGGAAT
TTTTCAAATAA

*Ppcaomt* (His$_6$-tagged)

ATGGGCAGCAGCCATCATCATCATCATCACAGCAGCGGCCTGGTGCCGCGCGGCAGCCATATGGCAAGCAGCCTGGAACGTAAAAGCC
ATCCGAAAATTAACCATGCAGAACCGGAAGATGAAATCACCAAAGAAGAAGAGGACGAGAGCTTTTGTTATGCAATGCAGCTGGTTGG
TAGCAGCGTTCTGAGCATGAGCCTGCAGAGCGCAATTAAACTGGGTATCTTTGATATCATTGCACGTAAAGGTCCGGGTGCAAAACTG
AGCAGCAGCGAAATTGCAACCAAAATTGGCACCGAAAATCCGGAAGCACCGGTTATGGTTGATCGTATTCTGCGTCTGCTGACCAGCC
ATAGCGTGCTGAATTGTAGCGCAGTTGCAGCAAATGGTGGTAGCGATTTTCAGCGTGTTTATAGCCTGGGTCCTGTTAGCAAATATTT
CGTGAATGATGAAGAAGGTGGTAGTCTGGGTCCGCTGCTGACCCTGATTCAGGATCGTGTTTTTCTGGAAAGCTGGTCACAGCTGAAA
GATGCAGTTGTTGAAGGTGGCATTCCGTTTAATCGTGTTCATGGTATGCATGCCTTTGAATATCCTGGTCTGGATCCGCGTTTTAATC
AGGTGTTTAATACCGCCATGTTTAACCATACAACCATCGTGATCAAAAAGCTGCTGCATATCTATAAAGGCCTGGAAGATAAAAATCT
GACCCAGCTGGTAGATGTTGGTGGTGGCCTGGGTGTTACCCTGAATCTGATTACCAGCCGTTATCAGCATATCAAGGGCATTAACTTT
GATCTGCCGCATGTTGTTAATCATGCACCGAGCTATCCGGGTGTTGAACATGTTGGCGGTGATATGTTTGCAAGCGTTCCGAGCGGTG
ATGCCATTTTTATGAAATGGATTCTGCATGATTGGAGCGACGAACATTGTCTGAAACTGCTGAAAAATTGCTATAAAGCGATTCCGGA
TAACGGCAAAGTTATTGTTGTGGAAGCACTGCTGCCTGCAATGCCGGAAACCAGCACCGCCACCAAAACCACCAGTCAGCTGGATGTT
CTGATGATGACCCAGAATCCTGGTGGTAAAGAACGTAGCGAACAAGAATTTATGGCACTGGCAACCGGTGCAGGTTTTAGCGGTATTC
GTTATGAATGCTTTGTGTGCAATTTCTGGGTGATGGAATTTTTCAAATAA

*couO* (His$_6$-tagged)

ATGGGCAGCAGCCATCATCATCATCATCACAGCAGCGGCCTGGTGCCGCGCGGCAGCCATATGGCTGCCGCGCGGCACCAGGCCGCTG
CTGTGATGATGATGATGATGGCTGCTGCCCATATGAAAATTGAACCGATTACCGGCAGCGAAGCGGAAGCGTTTCATCGCATGGGCAG
CCGCGCGTTTGAACGCTATAACGAATTTGTGGATCTGCTGGTGGGCGCGGGCATTGCGGATGGCCAGACCGTGGTGGATCTGTGCTGC
GGCAGCGGCGAACTGGAAATTATTCTGACCAGCCGCTTTCCGAGCCTGAACCTGGTGGGCGTGGATCTGAGCGAAGATATGGTGCGCA
TTGCGCGCGATTATGCGGCGGAACAGGGCAAAGAACTGGAATTTCGCCATGGCGATGCGCAGAGCCCGGCGGGCATGGAAGATCTGCT
GGGCAAAGCGGATCTGGTGGTGAGCCGCCATGCGTTTCATCGCCTGACCCGCCTGCCGGCGGGCTTTGATACCATGCTGCGCCTGGTG
AAACCGGGCGGCGCGATTCTGAACGTGAGCTTTCTGCATCTGAGCGATTTTGATGAACCGGGCTTTCGCACCTGGGTGCGCTTTCTGA
AAGAACGCCCGTGGGATGCGGAAATGCAGGTGGCGTGGGCGCTGGCGCATTATTATGCGCCGCGCCTGCAGGATTATCGCGATGCGCT
GGCGCAGGCGGCGGATGAAACCCCGGTGAGCGAACAGCGCATTTGGGTGGATGATCAGGGCTATGGCGTGGCGACCGTGAAATGCTTT
GCGCGCCGCGCGGCGGCG

*ecmat* (His$_6$-tagged)

ATGGCAAAACACCTTTTTACGTCCGAGTCCGTCTCTGAAGGGCATCCTGACAAAATTGCTGACCAAATCTCTGATGCCGTTTTAGACG
CGATCCTCGAACAGGATCCGAAAGCACGCGTTGCTTGCGAAACCTACGTAAAAACCGGCATGGTTTTAGTTGGCGGCGAAATCACCAC
CAGCGCCTGGGTAGACATCGAAGAGATCACCCGTAACACCGTTCGCGAAATTGGCTATGTGCATTCCGACATGGGCTTTGACGCTAAC
TCCTGTGCAGTTCTGAGCGCTATCGGCAAACAGTCTCCTGACATCAACCAGGGCGTTGACCGTGCCGATCCGCTGGAACAGGGCGCGG
GTGACCAGGGTCTGATGTTTGGCTACGCAACTAATGAAACCGACGTGCTGATGCCAGCACCTATCACCTATGCACACCGTCTGGTACA
GCGTCAGGCTGAAGTGCGTAAAAACGGCACTCTGCCGTGGCTGCGCCCGGACGCGAAAAGCCAGGTGACTTTCCAGTATGACGACGGC
AAAATCGTTGGTATCGATGCTGTCGTGCTTTCCACTCAGCACTCTGAAGAGATCGACCAGAAATCGCTGCAAGAAGCGGTAATGGAAG
AGATCATCAAGCCGATTCTGCCCGCTGAATGGCTGACTTCTGCCACCAAATTCTTCATCAACCCGACCGGTCGTTTTGTTATCGGTGG
CCCGATGGGTGACTGCGGTCTGACTGGTCGTAAAATTATCGTTGATACCTACGGCGGCATGGCGCGTCACGGTGGCGGTGCATTCTCT
GGTAAAGATCCATCAAAAGTGGACCGTTCCGCAGCCTACGCAGCACGTTATGTCGCGAAAAACATCGTTGCTGCTGGCCTGGCCGATC
GTTGTGAAATTCAGGTTTCCTACGCAATCGGCGTGGCTGAACCGACTTCCATCATGGTAGAAACTTTCGGTACTGAGAAAGTGCCTTC
TGAACAACTGACTCTGCTGGTACGTGAGTTCTTCGACCTGCGCCCATACGGTCTGATTCAGATGCTGGATCTGCTGCACCCGATCTAC
AAAGAAACCGCAGCATACGGTCACTTTGGTCGTGAACATTTCCCGTGGGAAAAAACCGACAAAGCGCAGCTGCTGCGCGATGCTGCCG
GTCTGAAGTAA

*ecmtan* (His$_6$-tagged)

ATGGCTGCCGCGCGGCACCAGGCCGCTGCTGTGATGATGATGATGATGGCTGCTGCCCATATGAAAATCGGCATCATTGGTGCAATGG
AAGAAGAAGTTACGCTGCTGCGTGACAAAATCGAAAACCGTCAAACTATCAGTCTCGGCGGTTGCGAAATCTATACCGGCCAACTGAA
TGGAACCGAGGTTGCGCTTCTGAAATCGGGCATGGTAAAGTCGCTGCGGCGCTGGGTGCCACTTTGCTGTTGGAACACTGCAAGCCAG
ATGTGATTATTAACACCGGTTCTGCCGGTGGCCTGGCACCAACGTTGAAAGTGGGCGATATCGTTGTCTCGGACGAAGCACGTTATCA
CGACGCGGATGTCACGGCATTTGGTTATGAATACGGTCAGTTACCAGGCTGTCCGGCAGGCTTTAAAGCTGACGATAAACTGATCGCT
GCCGCTGAGGCCTGCATTGCCGAACTGAATCTTAACGCTGTACGTGGCCTGATTGTTAGCGGCGACGCTTTCATCAACGGTTCTGTTG
GTCTGGCGAAAATCCGCCACAACTTCCCACAGGCCATTGCTGTAGAGATGGAAGCGACGGCAATCGCCCATGTCTGCCACAATTTCAA
CGTCCCGTTTGTTGTCGTACGCGCCATCTCCGACGTGGCCGATCAACAGTCTCATCTTAGCTTCGATGAGTTCCTGGCTGTTGCCGCT
AAACAGTCCAGCCTGATGGTTGAGTCACTGGTGCAGAAACTTGCACATGGCTAA

*ecluxS* (His$_6$-tagged)

ATGGCTGCCGCGCGGCACCAGGCCGCTGCTGTGATGATGATGATGATGGCTGCTGCCCATATGCCGTTGTTAGATAGCTTCACAGTCG
ATCATACCCGGATGGAAGCGCCTGCAGTTCGGGTGGCGAAAACAATGAACACCCCGCATGGCGACGCAATCACCGTGTTCGATCTGCG
CTTCTGCGTGCCGAACAAAGAAGTGATGCCAGAAAGAGGGATCCATACCCTGGAGCACCTGTTTGCTGGTTTTATGCGTAACCATCTT
AACGGTAATGGTGTAGAGATTATCGATATCTCGCCAATGGGCTGCCGCACCGGTTTTTATATGAGTCTGATTGGTACGCCAGATGAGC
AGCGTGTTGCTGATGCCTGGAAAGCGGCAATGGAAGACGTGCTGAAAGTGCAGGATCAGAATCAGATCCCGGAACTGAACGTCTACCA
GTGTGGCACTTACCAGATGCACTCGTTGCAGGAAGCGCAGGATATTGCGCGTAGCATTCTGGAACGTGACGTACGCATCAACAGCAAC
GAAGAACTGGCACTGCCGAAAGAGAAGTTGCAGGAACTGCACATCTAG

*ecmetf* (His$_6$-tagged)

ATGGCTGCCGCGCGGCACCAGGCCGCTGCTGTGATGATGATGATGATGGCTGCTGCCCATATGAGCTTTTTTCACGCCAGCCAGCGGG
ATGCCCTGAATCAGAGCCTGGCAGAAGTCCAGGGGCAGATTAACGTTTCGTTCGAGTTTTTCCCGCCGCGTACCAGTGAAATGGAGCA
GACCCTGTGGAACTCCATCGATCGCCTTAGCAGCCTGAAACCGAAGTTTGTATCGGTGACCTATGGCGCGAACTCCGGCGAGCGCGAC
CGTACGCACAGCATTATTAAAGGCATTAAAGATCGCACTGGTCTGGAAGCGGCACCGCATCTTACTTGCATTGATGCGACGCCCGACG
AGCTGCGCACCATTGCACGCGACTACTGGAATAACGGTATTCGTCATATCGTGGCGCTGCGTGGCGATCTGCCGCCGGGAAGTGGTAA
GCCAGAAATGTATGCTTCTGACCTGGTGACGCTGTTAAAAGAAGTGGCAGATTTCGATATCTCCGTGGCGGCGTATCCGGAAGTTCAC
CCGGAAGCAAAAAGCGCTCAGGCGGATTTGCTTAATCTGAAACGCAAAGTGGATGCCGGAGCCAACCGCGCGATTACTCAGTTCTTCT
TCGATGTCGAAAGCTACCTGCGTTTTCGTGACCGCTGTGTATCGGCGGGCATTGATGTGGAAATTATTCCGGGAATTTTGCCGGTATC
TAACTTTAAACAGGCGAAGAAATTTGCCGATATGACCAACGTGCGTATTCCGGCGTGGATGGCGCAAATGTTCGACGGTCTGGATGAT
GATGCCGAAACCCGCAAACTGGTTGGCGCGAATATTGCCATGGATATGGTGAAGATTTTAAGCCGTGAAGGAGTGAAAGATTTCCACT
TCTATACGCTTAACCGTGCTGAAATGAGTTACGCGATTTGCCATACGCTGGGGGTTCGACCTGGTTTATAA

*mmsahh* (His$_6$-tagged)

ATGGCTGCCGCGCGGCACCAGGCCGCTGCTGTGATGATGATGATGATGGCTGCTGCCCATATGAGCGACAAACTGCCGTATAAAGTTG
CAGATATTGGTCTGGCAGCATGGGGTCGTAAAGCACTGGATATTGCAGAAAATGAAATGCCTGGTCTGATGCGTATGCGTGAAATGTA
TAGCGCAAGCAAACCGCTGAAAGGTGCACGTATTGCAGGTTGTCTGCACATGACCGTTGAAACCGCAGTTCTGATTGAAACCCTGGTT
GCACTGGGTGCAGAAGTTCGTTGGAGCAGCTGTAACATTTTTAGCACCCAGGATCATGCAGCAGCAGCAATTGCAAAAGCAGGTATTC
CGGTTTTTGCATGGAAAGGTGAAACCGATGAAGAATATCTGTGGTGTATTGAACAGACCCTGCACTTTAAAGATGGTCCGCTGAATAT
GATTCTGGATGATGGTGGTGATCTGACCAATCTGATTCATACCAAATATCCGCAGCTGCTGAGCGGTATTCGTGGTATTAGCGAAGAA
ACCACCACCGGTGTTCATAACCTGTATAAAATGATGAGCAACGGCATTCTGAAAGTTCCGGCAATTAATGTTAATGATAGCGTGACCA
AAAGCAAATTCGATAATCTGTATGGTTGTCGCGAAAGCCTGATTGATGGTATTAAACGTGCAACCGATGTTATGATTGCAGGTAAAGT
TGCCGTTGTTGCAGGTTATGGTGATGTTGGTAAAGGTTGTGCACAGGCACTGCGTGGTTTTGGTGCACGTGTGATTATTACCGAAATT
GATCCGATTAATGCACTGCAGGCAGCAATGGAAGGCTATGAAGTTACCACCATGGATGAAGCATGTAAAGAAGGCAACATTTTTGTTA
CAACCACCGGTTGCGTTGATATCATTCTGGGTCGTCATTTTGAGCAGATGAAAGATGATGCCATTGTGTGCAATATCGGCCATTTTGA
TGTTGAGATTGATGTGAAATGGCTGAATGAAAACGCCGTGGAAAAAGTGAACATTAAACCGCAGGTTGATCGCTATTGGCTGAAAAAT
GGTCGTCGTATTATTCTGCTGGCAGAAGGTCGTCTGGTTAATCTGGGTTGTGCAATGGGTCATCCGAGCTTTGTTATGAGCAATAGCT
TTACCAATCAGGTGATGGCACAGATTGAACTGTGGACCCATCCGGATAAATATCCGGTTGGTGTTCATTTCCTGCCGAAAAAACTGGA
TGAGGCAGTTGCAGAAGCACATCTGGGTAAACTGAATGTGAAACTGACCAAACTGACAGAAAAACAGGCACAGTATCTGGGTATGCCG
ATTAACGGTCCGTTTAAACCGGATCATTATCGCTATTAA
